# Simulating metagenomic stable isotope probing datasets with MetaSIPSim

**DOI:** 10.1101/735787

**Authors:** Samuel E. Barnett, Daniel H. Buckley

## Abstract

**Background:** DNA-stable isotope probing (DNA-SIP) links microorganisms to their *in-situ* function in diverse environmental samples. Combining DNA-SIP and metagenomics (metagenomic-SIP) allows us to link genomes from complex communities to their specific functions and improves the assembly and binning of these targeted genomes. However, empirical development of metagenomic-SIP methods is hindered by the complexity and cost of these studies. We developed a toolkit, ‘MetaSIPSim,’ to simulate sequencing read libraries for metagenomic-SIP experiments. MetaSIPSim is intended to generate datasets for method development and testing. To this end, we used MetaSIPSim generated data to demonstrate the advantages of metagenomic-SIP over a conventional shotgun metagenomic sequencing experiment.

**Results:** Through simulation we show that metagenomic-SIP improves the assembly and binning of isotopically labeled genomes relative to a conventional metagenomic approach. Improvements were dependent on experimental parameters and on sequencing depth. Community level G+C content impacted the assembly of labeled genomes and subsequent binning, where high community G+C generally reduced the benefits of metagenomic-SIP. Furthermore, when a high proportion of the community is isotopically labeled, the benefits of metagenomic-SIP decline. Finally, the choice of gradient fractions to sequence greatly influences method performance.

**Conclusions:** Metagenomic-SIP is a valuable method for recovering isotopically labeled genomes from complex communities. We show that metagenomic-SIP performance depends on optimization of experimental parameters. MetaSIPSim allows for simulation of metagenomic-SIP datasets which facilitates the optimization and development of metagenomic-SIP experiments and analytical approaches for dealing with these data.

## Introduction

DNA-Stable isotope probing (DNA-SIP) is a powerful tool for linking uncultured microorganisms to their function within environmental samples [1]. DNA-SIP has applications in a wide range of areas including biogeochemistry [2–9], biodegradation [10–14], and ecological interactions [15, 16]. In the past few years, methods such as high-resolution SIP (HR-SIP) [2, 17] and quantitative SIP (qSIP) [18], have been developed to analyze amplicon sequencing data from DNA-SIP experiments. DNA-SIP has also been combined with metagenomic sequencing (metagenomic-SIP) to link *in situ* metabolic activity to genome composition [13, 19, 20]. Metagenomic-SIP is believed to improve the recovery of metagenome-assembled genomes (MAGs) from ^13^C-labeled organisms [21, 22]. Unfortunately, validation and improvement of metagenomic-SIP methods and analytical tools have largely been hindered by the difficulty and cost of these experiments [14, 21].

A recently developed open-source toolkit, SIPSim [17], enables *in-silico* simulation of amplicon sequencing data from DNA-SIP experiments (*i.e.* species abundance tables). SIPSim has been used to compare various DNA-SIP analysis methods [17] and is useful for testing the design of DNA-SIP experiments, but does not simulate sequencing read libraries and therefore is of limited use for development and testing of metagenomic-SIP methods. From here on, we will use metagenomic-SIP to refer to shotgun metagenomic sequencing from a DNA-SIP experiment. Here, we present a newly developed toolkit, MetaSIPSim, to simulate metagenomic-SIP datasets. MetaSIPSim generates sequencing read libraries in FASTA or FASTQ format such as those generated in a metagenomic-SIP experiment. MetaSIPSim is freely available on Github (https://github.com/seb369/MetaSIPSim). At this time, there are no other tools publicly available that generate simulated metagenomic-SIP read libraries. Our tool will allow researchers to rapidly and inexpensively test experimental parameters and develop analytical tools which will advance the sophistication and accuracy of metagenomic-SIP experiments. SIP approaches are increasingly popular in environmental microbiology.

We demonstrated the utility of MetaSIPSim by assessing whether the coverage, assembly, and MAG binning of target ^13^C-labeled genomes were improved by metagenomic-SIP relative to a conventional shotgun metagenomic approach. We hypothesized that the enhanced performance of metagenomic-SIP relative to shotgun metagenomics depends on sequencing depth (*i.e.* number of reads recovered). To test this hypothesis, we ran all simulations at both 5,000,000 reads (5M) and 10,000,000 reads (10M). We predicted that the benefits of metagenomic-SIP for assembly and binning would be greatest when sequencing depth is lower (*i.e*. in 5M). Furthermore, we hypothesized that the performance of metagenomic-SIP varies with the guanine and cytosine (G+C) content of whole community DNA. G+C content determines where a DNA fragment localizes in a buoyant density (BD) gradient [23, 24]. Variation in individual genome G+C in complex communities has been shown to affect measurements of isotope incorporation in DNA-SIP experiments, such that unlabeled high G+C DNA can potentially co-migrate with labeled DNA of low G+C content [17, 25]. In addition, genomic G+C content is not uniform within individual genomes and the average G+C content of a genome fragment will differ from the average G+C content of its source genome with variance a function of fragment length. It is unclear the extent to which variation in community level G+C content, and intra-genomic variation in G+C content, affects metagenome-SIP analyses. To test the hypothesis that community-level genomic G+C content affects metagenomics-SIP performance, we simulated metagenomic-SIP experiments using communities that differ in G+C distribution (low, medium, and high). We predicted that read coverage, metagenome assembly and MAG bin quality would vary with community G+C content resulting in lower performance for high G+C communities.

## Materials and methods

### Implementation

MetaSIPSim was developed using the Ubuntu 16.04.4 operating system running python 2.7 and has been successfully run on Mac OSX 10.12.6. All dependencies and their development versions are provided in Table S1. MetaSIPSim can run with parallel processes to reduce running time. MetaSIPSim can be memory intensive, depending mostly on the number of reference sequences, reference sequence size, and number of reads generated. It is recommended to do a test run to make sure that the local system has enough RAM to handle a desired simulation.

### Simulation procedure

The input to MetaSIPSim is a configuration file with all parameters discussed below as well as input and output paths. One input is the directory containing the reference sequences. All reference sequences must be in FASTA format. A reference can be a whole genome, scaffolds, or contigs, but each reference must be in a separate FASTA file. If a reference is composed of multiple scaffolds, chromosomes, or plasmids, its file should be in multi-FASTA format. All scaffolds within this file will be processed as a single reference. A diagram of the simulation procedure is found in Fig. S1. The first step of the simulation is to fragment each reference into discrete sequence segments termed ‘fragments.’ Fragment size is based on a user provided distribution. The fragmentation process simulates DNA fragmentation patterns occurring during DNA extraction, such as from bead beating. Fragment size distributions can be of uniform, normal, truncated-normal, or skewed-normal distribution, which should be chosen based on empirical evidence from a user’s own extraction methods. The fragmentation process is repeated several times such that each reference has a fragment coverage designated by the user to get a diverse sample of fragments for each reference.

MetaSIPSim has the capacity to perform two simulation modes. The first simulation mode is the ‘single BD window method,’ similar to heavy-SIP [17], sequencing a single gradient window using BD boundaries (*ρ*_*max*_, *ρ*_*max*_) defined by the user. Most published metagenomic-SIP studies to date have employed this ‘heavy-SIP method.’ The second simulation mode treats each density gradient fraction independently, similar to HR-SIP [2], in which multiple fractions spanning the gradient are sequenced individually. For this fraction-based simulation mode, the simulation is performed independently for each fraction, with the individual fraction BD boundaries defining the gradient window (*ρ*_*max*_, *ρ*_*max*_). The fraction-based simulation mode uses the same reference fragments in all individual simulations. All variables used in the following equations are summarized in Table 1.

**Table 1:** Descriptions of variables used in simulation equations.

| Variable | Description |
| --- | --- |
| $G + C$ | Proportion of DNA that is guanine or cytosine |
| $\rho_t$ | Theoretical BD of a fragment based on fragment G+C |
| $\rho$ | BD of fragment after accounting for isotopic labeling |
| $\sigma$ | Standard deviation of fragment BD ($\rho$) |
| $\rho_{min}$ | Minimum BD of window to be sequenced |
| $\rho_{max}$ | Maximum BD of window to be sequenced |
| $\rho_{DBLmin}$ | Minimum BD of the DBL for a fragment |
| $\rho_{DBLmax}$ | Maximum BD of the DBL for a fragment |
| $A$ | Atom % excess of the fragment |
| $\delta$ | Increase in DNA BD if atom % excess is 100% ($^{13}\text{C}$: 0.036, $^{15}\text{N}$: 0.016) |
| $p_{DBL}$ | Proportion of DNA found in the diffusive boundary layer. $1 - p_{DBL}$ is the proportion of DNA found in the lumen of the tube |
| $p_{LR}$ | Proportion of the fragment lumen population recovered in the BD window |
| $r_t$ | Radius of the ultracentrifuge tube |
| $r_{min}$ | Minimum distance from the axis of rotation to the tube |
| $r_{max}$ | Maximum distance from the axis of rotation to the tube |
| $x$ | Fragment distance from the axis of rotation at equilibrium |
| $x_{min}$ | Minimum position on tube of the DBL range for a fragment |
| $x_{max}$ | Maximum position on tube of the DBL range for a fragment |
| $\alpha$ | Abundance of the genome/fragment in the sample |
| $\alpha_L$ | Abundance of the fragment in the lumen of the tube within the BD |
| $\alpha_{DBL}$ | Abundance of a fragment in the DBL within the BD window |
| $\alpha_f$ | Abundance of the fragment in the BD window |
| $\alpha_r$ | Abundance of reads from the fragment |
| $l$ | Length of the fragment in base pairs |
| $l_r$ | Read length in base pairs |
| $R$ | Universal gas constant (8.314 J/molK) |
| $T$ | Temperature in kelvin |
| $\beta$ | Proportionality constant of aqueous cesium chloride ($1.14 \times 10^9$) |
| $G$ | Buoyancy factor ($7.87 \times 10^{-10}$) |
| $M_c$ | Mean molar weight of standard nucleotide base pair in cesium chloride solution (882) |
| $D$ | Average density of the gradient solution |
| $\omega$ | Angular velocity of centrifugation |
| $I$ | Isoconcentration point |
| $\theta$ | Angle of the tube relative to the axis of rotation in radians |

Following reference sequence fragmentation, the abundance of each fragment within the gradient window is estimated. The initial fragment abundance (*α*) is equal to the relative abundance of the parent reference, provided in a user supplied community composition table. The abundance of each fragment in the gradient window is then determined as a function of fragment BD characteristics. The theoretical BD for the fragment (*ρ*_*t*_) is calculated from the *G* + *C* of the fragment (Equation 1) [26]:

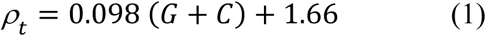

The theoretical BD is then adjusted for isotopic labeling based on the atom % excess (*A*) assigned to parent reference. The atom % excess for an isotopically labeled fragment is randomly generated from a normal distribution, with the mean and standard deviation for each parent reference supplied in a user supplied incorporator identification table. With this setup, users can set different incorporators to varying levels of isotope labeling. If the fragment is from an unlabeled reference, the atom % excess is 0. The mean BD (*ρ*) of each fragment is calculated as such (Equation 2) [17]:

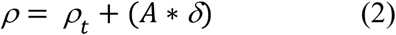

Here, *δ* is the increase in BD of DNA if the atom % excess is 100% (*i.e.* if the DNA was fully isotopically labeled). *δ* differs by isotope where *δ* = 0.036 for ^13^C and *δ* = 0.016 for ^15^N [24].

Next, it is necessary to calculate ‘DNA smearing’ due to diffusive boundary layer effects during the deceleration stage of ultracentrifugation when gradient reorientation occurs (Fig. S2) [17]. Most DNA is present in the ‘lumen’ of the centrifuge tube, which is defined as all DNA distant from and not interacting with the ultracentrifuge tube walls (Fig. S2). When the gradient reorients during deceleration, interactions with tube walls causes a thin layer of gradient solution containing DNA to be trapped within the diffusive boundary layer (DBL) along the tube walls [17]. The DBL does not move with the lumen-DNA during reorientation and, during fractionation, this unequilibrated ‘DBL-DNA’ will contaminate fractions with which it intersects (Fig. S2). The proportion of DNA in the DBL is minor compared to the lumen DNA, but is readily detected with high throughput sequencing approaches [17]. For simplicity in the simulation, the proportion of DNA found in the DBL (*p*_*DBL*_) relative to the total DNA concentration is provided by the user. Empirical studies are needed to determine the equations and experimental properties governing the ratio of DBL-DNA to lumen DNA.

Within the tube lumen, due to diffusion, a fragment will be normally distributed around the isotope adjusted BD (*ρ*) and standard deviation (*σ*, Equation 3; Fig. S2) [27]:

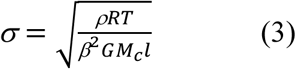

Where *R* is the universal gas constant (8.314 J/molK), *T* is the temperature in Kelvin, *β* is the proportionality constant of aqueous cesium chloride (1.14 × 10^9^)[28], *G* is a buoyancy factor (7.87 × 10^−10^)[29], *M*_C_ is the mean molecular weight of a standard nucleotide base pair in cesium chloride solution (882 g/mol)[29], and *l* is the length of the fragment in base pairs. The proportion of the fragment lumen-population recovered in the BD window (*p*_*LR*_) is then calculated from the cumulative density function of the normal distribution with mean *ρ* and standard deviation *σ* and bounded by the maximum and minimum buoyant densities of the gradient window (*ρ*_*max*_, *ρ*_*max*_; Fig. S2). Thus, the abundance of the lumen-fragment (α_*L*_) in the window is the total lumen abundance multiplied by this proportion recovered in the window (Equation 4).

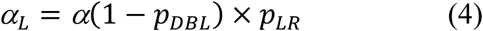

To calculate the abundance of the DBL-fragment recovered in the gradient window, the range of buoyant densities that the DBL-fragment is contaminating is determined. First, the distance from the axis of rotation that the fragment will be found (*x*) at equilibrium is calculated (Equation 5)[24]:

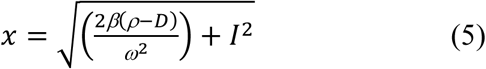

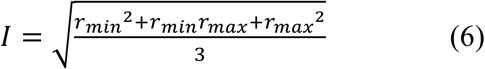

Here, *D* is the average density of the gradient in g/ml, *ω* is the angular velocity of centrifugation in rad/s, and *I* is the isoconcentration point (Equation 6 [24]). *r*_*min*_ and *r*_*max*_ are the minimum and maximum distances from the axis of rotation to the tube (Fig. S2). From this, the minimum position (*x*_*min*_) of the DBL range along the ultracentrifuge tube can be calculated both if found in the middle, cylindrical section (Equation 7) or the bottom, rounded section (Equation 8) of the ultracentrifuge tube (Fig. S2)[17].

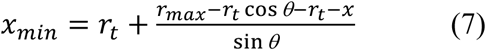

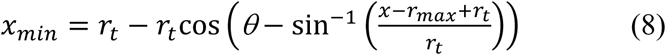

Here, *θ* is the angle of the tube relative to the axis of rotation in radians, *r*_*t*_ is the radius of the ultracentrifuge tube. Similarly, the maximum positions (*x*_*max*_) of the DBL range is calculated whether in the middle, cylindrical section (Equation 9) or the bottom, rounded section (Equation 10) of the ultracentrifuge tube [17].

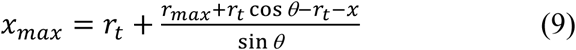

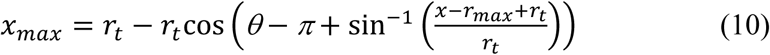

Then the maximum and minimum positions are converted into BD limits (*ρ*_*DBLmin*_and *ρ*_*DBLmax*_) from a table generated with a model gradient. Model gradients are generated as in SIPSim [17].

The abundance of the DBL-fragment (α_*DBL*_) recovered in the gradient window is the proportion of the DBL BD range covered by the window then multiplied by the total abundance of the fragment DBL-population (Equation 11).

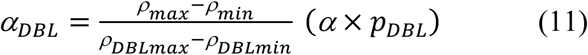

From the previous calculations, the abundance of a fragment recovered in the sequenced BD window (α_*f*_) is simply the sum of the abundances of lumen and DBL fragments (Equation 12).

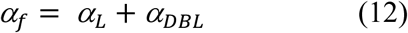

Next, sequencing reads are generated from the fragments. To do this, the estimated relative abundance of each potential read (α_*r*_) derived from the fragment is determined (Equation 13) based on the lengths of the fragment (*l*_*f*_) and of the reads (*l*_*r*_).

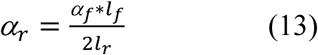

Common read lengths for Illumina sequencing chemistry include 125, 150, 250, and 300 bp. The initial read abundance is transformed into a relative abundance by dividing by the sum of all read abundances across the fragment. Then the number of reads to be recovered from each fragment is assigned randomly, weighted by the relative abundance of reads within each fragment. Each paired end read from each fragment is then generated by randomly selecting forward and reverse read starting points based on the read length and insert size. Insert size is randomly generated for each read based on a normal distribution with mean and standard deviation provided by the user. Forward or reverse assignment is random. Finally, read sequences are retrieved from the original reference sequence based on coordinates which have been propagated from fragment to sequencing read. Reverse reads are converted to the reverse compliment. All final forward and reverse read sequences are written out to two multi-FASTA files with paired unique identifiers.

To compare metagenomic-SIP data to a standard shotgun metagenomic dataset, MetaSIPSim includes a script for generating bulk community metagenomes with the same references and parameters as the metagenomic-SIP simulation. This script functions similarly to the metagenomic-SIP simulator, using the same input configuration file. However, in this case the abundance of each simulated fragment (α_*f*_) is equal to the reference abundance (*α*), without adjustments for gradient fractionation. The FASTA output generated by MetaSIPSim has no sequencing errors or quality score information associated with high throughput sequencing datasets. We have included a script that includes modified functions from InSilicoSeq [30] to convert the FASTA file output into FASTQ format by simulating sequencing errors and quality scores. The conversion uses an error model included in the InSilicoSeq installation or created by the user. The error models are sequencing platform specific, with options for Illumina MiSeq, HiSeq, or NovaSeq. Unlike the original InSilicoSeq implementation, the MetaSIPSim script does not add gaps to sequences.

### Validation

We validated the ability of MetaSIPSim to simulate the distribution of both isotopically labeled and unlabeled genomic DNA across a CsCl gradient against three published DNA-SIP studies [25, 31, 32]. These studies used bacterial isolates grown on ^13^C-labeled, ^15^N-labeled, or unlabeled substrates (Table 2). For each study, we simulated fragments across multiple CsCl gradient fractions. Fraction BD boundaries were estimated based on the reported average fraction BD. Gradient parameters were taken from those reported (Table S2). We used a uniform atom % excess of 100% as these studies used pure cultures and fully labeled susbtrates. The code for validation simulations available at https://github.com/seb369/MetaSIPSim/validation/.

**Table 2:**
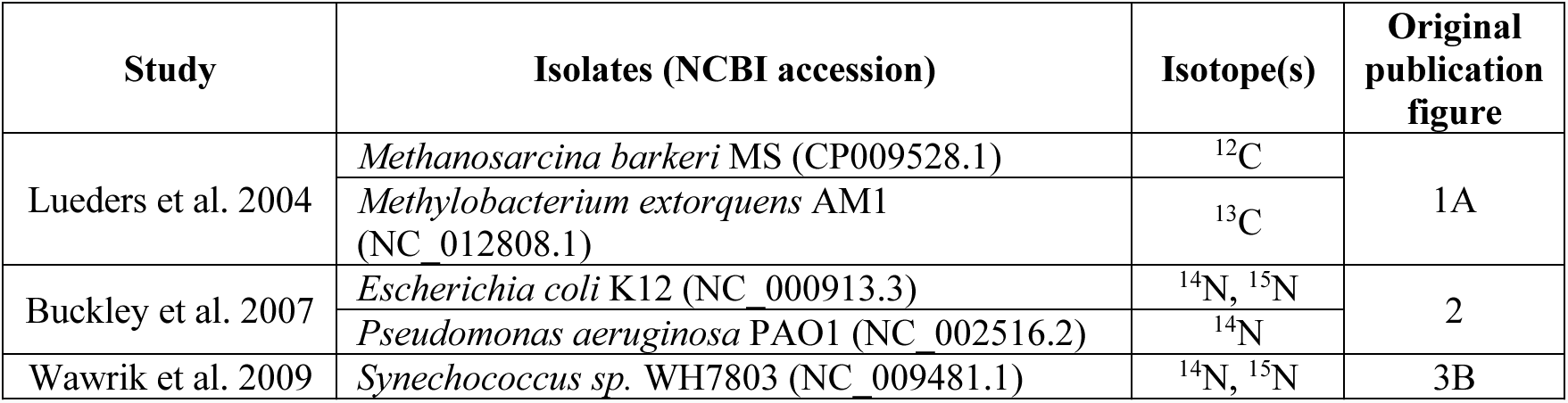
Studies used for validation of fragment abundance distributions including the isolates reported in the study, NCBI accessions used to download the genomes, and the stable isotopes simulated

| Study | Isolates (NCBI accession) | Isotope(s) | Original publication figure |
| --- | --- | --- | --- |
| Lueders et al. 2004 | *Methanosarcina barkeri* MS (CP009528.1) | <sup>12</sup>C | 1A |
|  | *Methylobacterium extorquens* AM1 (NC_012808.1) | <sup>13</sup>C |  |
| Buckley et al. 2007 | *Escherichia coli* K12 (NC_000913.3) | <sup>14</sup>N, <sup>15</sup>N | 2 |
|  | *Pseudomonas aeruginosa* PAO1 (NC_002516.2) | <sup>14</sup>N |  |
| Wawrik et al. 2009 | *Synechococcus sp.* WH7803 (NC_009481.1) | <sup>14</sup>N, <sup>15</sup>N | 3B |

All three studies measured the amount of genomic DNA of an isolate recovered across fractions, normalized to the fraction with the highest DNA concentration. To approximate this value, within each fraction, we multiplied the abundance of each fragment by its length, then summed this base pair count across all fragments get a quantity of genomic DNA. We then normalized to the fraction with the greatest quantity of DNA to get the gradient profile. Empirical distributions were estimated from the published figures using Engage Digitizer version 6.0 [33]. We ran simulations with SIPSim [17] using the same parameters as an additional comparison.

### Case study: Assessment of improved metagenomes using metagenomic-SIP

We used MetaSIPSim to assess whether metagenomic-SIP improves coverage, assembly, and binning of labeled bacterial genomes compared to conventional shotgun metagenome sequencing. The simulated SIP experiment was based on a ^13^C, heavy window metagenomic-SIP experiment. A diagram of the experimental design is found in Fig. S3. A total of 1542 reference genomes were downloaded from the NCBI RefSeq database [34] on January 25, 2019. Genomes that were related at the species level, based on reported taxonomy, were pruned such that a single genome was present per species. The remaining 1491 reference genomes had a bimodal distribution of G+C, with peaks around 40 and 65% (Fig. S4). To get a representative set of reference genomes with a relatively uniform G+C distribution for sampling, we sampled 20 genomes from each integer between 20-75% G+C after rounding G+C values to the nearest integer. If 20 or fewer genomes were present at a given G+C integer, then all genomes with that G+C value were selected. From this subset of genomes, we then randomly subsetted 500 genomes to meet G+C content criteria for the reference sets: lowGC (40%), medGC (50%), and highGC (60%). Genome selection was weighted by the absolute value of the distance of each genome’s G+C to the target community G+C (Fig. S5).

From each reference set, we generated six replicate synthetic communities with randomized species abundance distributions. These replicate communities represent the variation one would expect from a SIP experiment with biological replicates, such as independent microcosms. The compositions of each replicate community was generated using the *communities* function from SIPSim [17] with all genomes present (richness = 1) in each replicate, with a lognormal distribution, a mean relative abundance of 2.0%, and standard deviation of 0.8. All genomes from the reference set were found in each community but 10% of abundance ranks were permuted, to provide more realistic variation between communities. Overall, we simulated a total of 18 communities. The first replicate community from each reference set was designated as the control, in which no genomes were isotopically labeled.

We randomly selected 20% of each reference set (*i.e.* 100 genomes) to be isotopically enriched ‘incorporators.’ To avoid a reference set with incorporators disproportionality weighted to either high or low abundance compared to the other sets, genomes were selected such that the mean abundance ranks of incorporators were similar across all three reference sets (Fig S6). For each treatment community, 50 genomes from the incorporator set were randomly chosen to be labeled. This process was reiterated until all 100 incorporators were assigned to at least one treatment community. All incorporators had a mean atom % excess of ^13^C of 90% and standard deviation of 5. This experimental design, with five treatment and one control sample, where labeled genomes may vary between treatment samples represents a metagenomic-SIP experiment where different labeled substrates were supplied to enrich different populations within a community [2, 35]. In this type of experiment organisms can be labeled under more than one treatment and one unlabeled replicate community can act as a control for multiple treatments. CsCl gradients were simulated for each sample using the *gradient_fractions* function from SIPSim [17] with minimum and maximum buoyant densities of 1.675 and 1.771 g/ml.

We simulated both metagenomic-SIP and conventional shotgun metagenome reads from the synthetic communities using MetaSIPSim. SIP gradient parameters were derived from Pepe-Ranney *et al*. [2] (Table S2). These parameters were used as they represent a standard DNA-SIP experiment used in a number of studies. For both types of simulations, we generated 5,000,000 (5M) and 10,000,000 (10M) paired end, 151 bp reads with an average insert size of 1,000 bp and standard deviation of 5. Sequencing errors and quality scores were added using the NovaSeq error model from InSilicoSeq [30]. We performed the following metagenome processing pipeline to assemble and bin contigs separately for each reference set, read depth, and metagenome simulation type: (*i*) co-assembly was performed with the six read libraries, one per replicate community, using MEGAHIT version 1.1.3 [36] with default parameters, and (*ii*) contigs were binned with MetaBAT2 version 2.12.1 [37, 38] using default parameters. MetaBAT2 was chosen as it incorporates differential abundance binning, which takes advantage of the differential labeling of genomes across treatments [22]. All code for these simulations including metagenome data processing are available at https://github.com/seb369/MetaSIPSim/case_study/.

We assessed how well each reference genome was recovered in raw reads by separately mapping libraries to the references using BBMap version 37.10 [39]. Specifically, we used the genome coverage and the proportion of the genome completely mapped by reads as indicators. We assessed assembly quality through alignment of contigs to the reference genomes using MetaQUAST version 5.0.2 [40]. The metrics used to gauge the assembly quality were the proportion of each reference genome aligned to contigs and the NGA50 for each reference. Successful MAG binning was assessed by first determining which genome was most aligned to each MAG. The proportion of the reference recovered as a MAG was calculated as length of the genome aligned to the binned contigs divided by genome length. Sections of the reference genome where multiple contigs aligned were only counted once (*i.e.* contig overlaps). Bin contamination was calculated as the summed length of contig regions not aligned to the assigned reference divided by the total bin size. In all analyses, we compared assessment metrics between the metagenomic-SIP and the shotgun metagenomic datasets. All statistical analyses were performed in R version 3.4.4 [41] using the Wilcoxon signed rank tests.

We ran two follow up analyses based on trends observed with initial simulations. First, we tested whether the number of isotopically labeled genomes within each sample influenced metagenomic-SIP performance. This simulation was done using the lowGC reference set. Incorporators were randomly selected as before, however we selected 25 incorporators per treatment (from 50 total incorporators in the reference set) or 100 incorporators per treatment (from 200 in the reference set). All other community and simulation parameters were identical to the original simulations. Secondly, we tested whether the selection of the BD window to sequence influences metagenomic-SIP improvement. These simulations were done using the highGC reference set. All community and simulation parameters were identical as before except that the BD window to be sequenced was 1.70-1.75 g/ml, 1.72-1.77 g/ml, or 1.75-1.79 g/ml. The BD range for the model gradients were also slightly extended to 1.67-1.80 g/ml to account for the new window ranges. We simulated 5,000,000 reads for all follow up simulations. All metagenome processing and analyses were performed as previously described.

## Results

### Implementation

Simulations for the metagenomic-SIP case study took on average 98 minutes and 121 minutes for six replicate communities at 5,000,000 and 10,000,000 reads respectively, using 10 processors. The corresponding simulations for conventional shotgun sequencing took approximately 38 minutes and 56 minutes respectively. Fragment generation took up to 10 minutes of those runs, with all additional time required for generating reads for each of the six replicate communities. These processing times do not include generation of input files, conversion from FASTA to FASTQ formats, or any read processing or analysis. Overall the simulations used less than 20 GB of RAM on an Ubuntu 16.04.4 operating system.

### Validation

We simulated the distribution of fragmented genomic DNA across CsCl gradients based on experimental procedures from three published studies (Table 2). We found that the peak genomic DNA density recovery from the MetaSIPSim and SIPSim simulations roughly matched the empirical results from Buckley et al. 2007 [25] and Wawrik et al. 2009 [32] (Fig. 1B and C) and the general distributions were similar across datasets for these studies. Variations in overall distributions are likely due in part to variability from experimental conditions and variations in genome composition between isolate and reference sequence strains. For the Lueders et al. 2004 [31] study, the peak genomic DNA density matched between MetaSIPSim and SIPSim, however these peaks were in adjacent BD fractions compared to the empirical data (Fig. 1A). This small difference may be due to variation in experimental parameters or methodology that were not simulated.

**Fig. 1:**
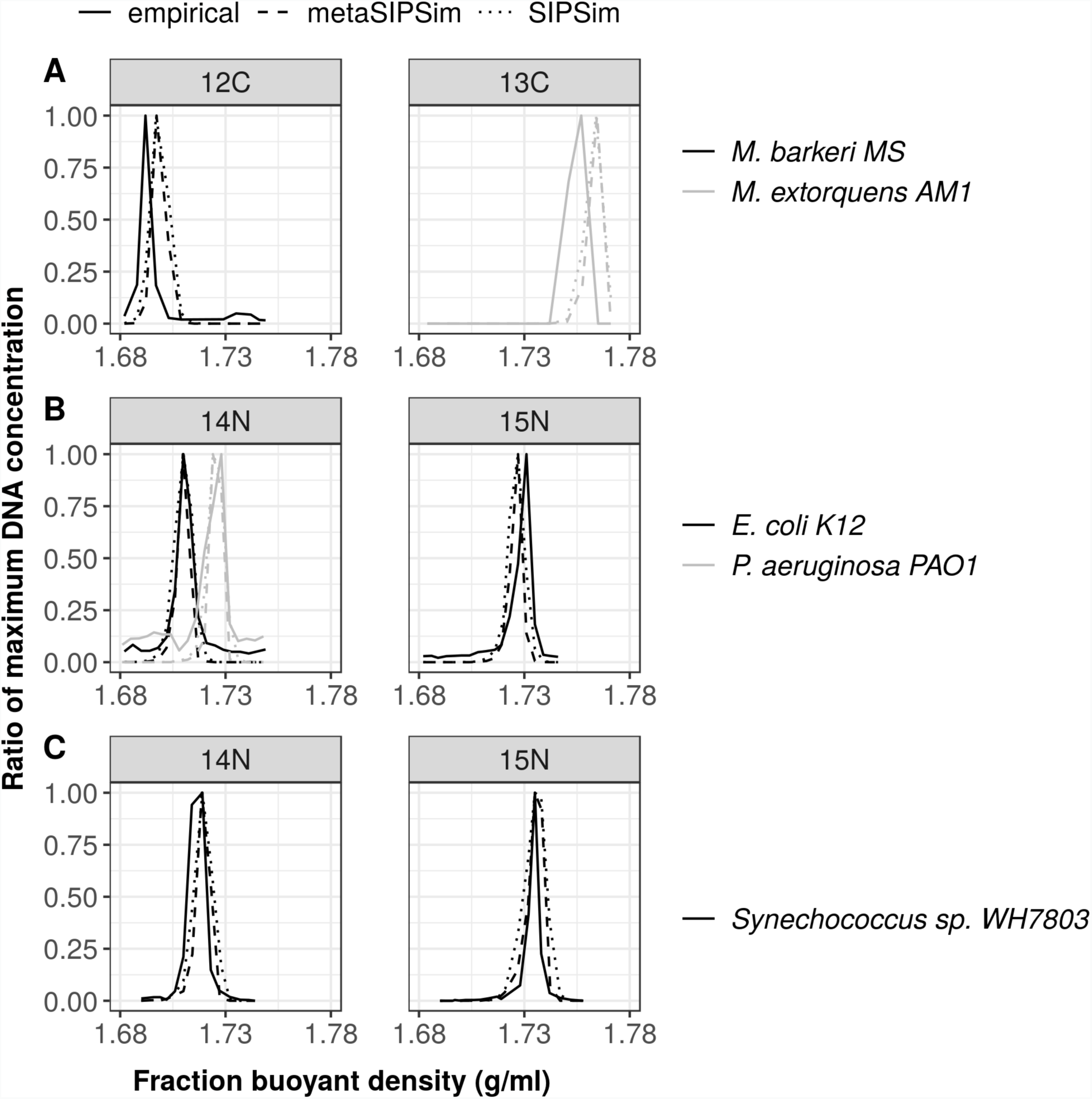
Comparisons among MetaSIPSim and SIPSim simulated genomic DNA distribution across CsCl gradients and the empirical results from A) Lueders et al. 2004 [31], B) Buckley et al. 2007 [25], and C) Wawrik et al. 2009 [32]

### Metagenomic-SIP improves recovery of isotopically labeled genomes in raw reads

We used MetaSIPSim to assess the ability of metagenomic-SIP to improve genome assembly and binning relative to shotgun metagenomics. Metagenomic-SIP should enrich for reads from isotopically labeled genomes (*i.e.* incorporators), so we first examined coverage and recovery of incorporator genomes in our raw simulated reads. As expected, we found that incorporators had greater coverage and were recovered more completely in the metagenomic-SIP simulation compared to paired conventional metagenomic libraries (Fig. 2A and C, Table 3). The difference in incorporator coverage and genome recovery between metagenomic-SIP and shotgun metagenomes were significantly greater than zero across all simulations (all *p*-values < 0.001; Table 3). The increased coverage achieved by metagenomic-SIP was negatively correlated with community G+C. The increase in genome recovery in raw reads with metagenomic-SIP was highest for the medGC reference set. The increase in coverage with metagenomic-SIP for each incorporator was also strongly affected by the G+C content of the target genome, where low G+C genomes had the least fold difference in coverage with metagenomic-SIP compared to shotgun metagenomics (Fig. S7). The improvement that metagenomic-SIP provides in the recovery of labeled genomes within raw reads was also somewhat affected by genome G+C but was more strongly driven by the relative abundance of target genomes in the community, with metagenomic-SIP providing the greatest benefit in recovering low abundance genomes (Fig. S8).

**Table 3:** Median difference 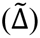 in metagenome quality for labeled genomes between simulations of metagenomic-SIP and conventional shotgun metagenomic data. For all measures, except bin contamination, a difference greater than zero indicates that metagenomic-SIP improved metagenome quality relative to the conventional metagenomics approach. For bin contamination, a difference less than zero indicates improved metagenome quality with metagenomic-SIP. All statistical analyses were single sided, Wilcoxon signed rank tests with an alternate hypothesis of greater than zero, except for bin contamination which used an alternate hypothesis of less than zero. All *p*-values are adjusted for multiple comparisons (Bonferroni, *n* = 6).

| Community G+C | Seq. depth | $\tilde{\Delta}$ Read coverage X (p-value) | $\tilde{\Delta}$ Proportion of genome recovered in reads (p-value) | $\tilde{\Delta}$ Proportion of genome recovered in contigs (p-value) | $\tilde{\Delta}$ NGA50 bp (p-value) | $\tilde{\Delta}$ Proportion of genome recovered in best bin (p-value) | $\tilde{\Delta}$ Proportion bin contaminated (p-value) |
| --- | --- | --- | --- | --- | --- | --- | --- |
| Low | 5M | 1.769<br>($< 0.001$) | 0.391<br>($< 0.001$) | 0.052<br>($< 0.001$) | 19672<br>($< 0.001$) | 0.567<br>($< 0.001$) | -0.038<br>($< 0.001$) |
| Medium | | 1.002<br>($< 0.001$) | 0.315<br>($< 0.001$) | 0.053<br>($< 0.001$) | 9251<br>($< 0.001$) | 0.353<br>($< 0.001$) | -0.022<br>($< 0.001$) |
| High | | 0.596<br>($< 0.001$) | 0.240<br>($< 0.001$) | 0.026<br>(NS) | 3511<br>($< 0.001$) | 0.366<br>($< 0.001$) | -0.010<br>($< 0.001$) |
| Low | 10M | 3.561<br>($< 0.001$) | 0.237<br>($< 0.001$) | 0.004<br>(0.007) | 11636<br>($< 0.001$) | 0.144<br>($< 0.001$) | -0.017<br>($< 0.001$) |
| Medium | | 1.992<br>($< 0.001$) | 0.306<br>($< 0.001$) | 0.007<br>($< 0.001$) | 7554<br>($< 0.001$) | 0.092<br>($< 0.001$) | -0.011<br>($< 0.001$) |
| High | | 1.174<br>($< 0.001$) | 0.222<br>($< 0.001$) | 0.001<br>(NS) | 7718<br>($< 0.001$) | 0.134<br>($< 0.001$) | -0.009<br>($< 0.001$) |

**Fig. 2:**
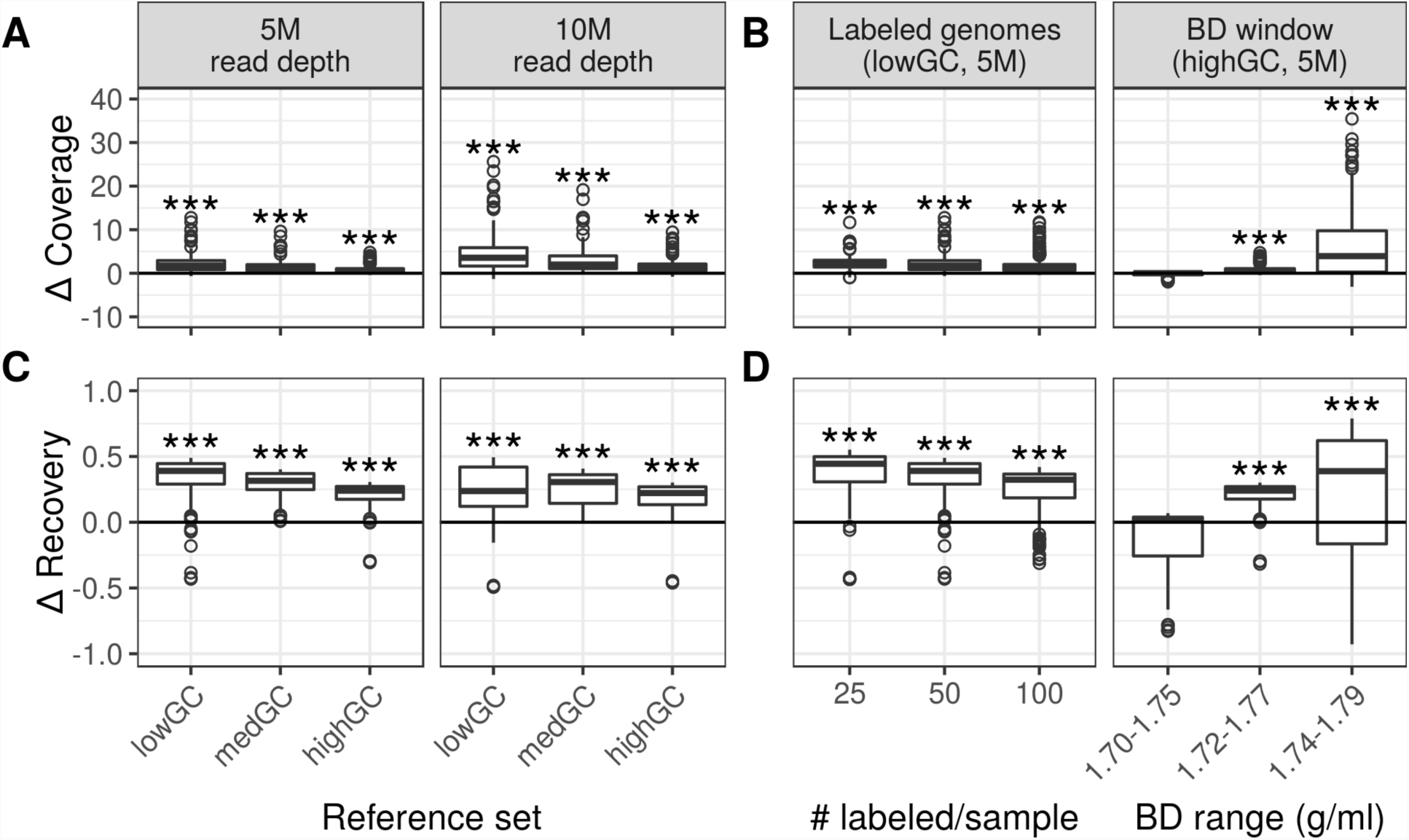
Improvement of metagenomic-SIP relative to conventional metagenomics with respect to coverage and recovery of ^13^C-labeled genomes with raw reads. Values greater than zero indicate improvement of metagenomic-SIP relative to conventional metagenomics (Bonferroni correction, *n* = 6; ***, *p*-values < 0.001). A and C) The difference in labeled genome (*n* = 100) coverage and the difference in proportion of the genome recovered (*i.e.* mapped to reads), respectively, varies between communities that differ in G+C content when sequenced at different depths. B and D) The difference in labeled genome coverage and the difference in proportion of the genome recovered (*i.e.* mapped to reads, *n* = 50, 100, 200), respectively, for low G+C communities with respect to the number of labeled genomes per sample and for high G+C communities with respect to the position of the BD window.

The number of incorporators had a minor impact on both labeled genome coverage and recovery within raw reads. The benefits of metagenomic-SIP relative to shotgun metagenomics were greatest when the number of incorporator genomes was low (Fig. 2B and D; Table 4). BD window position strongly influenced the coverage and recovery of targeted genomes within raw reads. Metagenomic-SIP analysis of a lighter BD window (1.70-1.75 g/ml) showed little improvement relative to standard metagenomics sequencing. In contrast, metagenomic-SIP analysis of a heavier windows improved significantly the coverage and recovery of target genomes within raw reads (1.72-1.77 g/ml and 1.74-1.79 g/ml; Fig. 2B and D; Table 4). For both measures, there was a strong relationship between the incorporator G+C and the BD of sequenced fractions. Metagenomic-SIP did not significantly improve coverage or recovery of raw reads from high G+C genomes in the light BD window over shotgun metagenomics (Fig. S9). Conversely, metagenomic-SIP did little to improve read coverage or recovery of low G+C genomes in the heavier BD window compared to the conventional approach (Fig. S10).

**Table 4:** Median difference 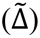 in metagenome quality measures for labeled genomes between simulations of metagenomic-SIP and conventional metagenomics. For all measures, except bin contamination, a difference greater than zero indicates that metagenomic-SIP improved metagenome quality relative to conventional metagenomics. For bin contamination, a difference less than zero indicates improved metagenome quality with metagenomic-SIP. All statistical analyses were single sided, Wilcoxon signed rank tests with an alternate hypothesis of greater than zero, except for bin contamination which used an alternate hypothesis of less than zero. All *p*-values are adjusted for multiple comparisons (Bonferroni, *n* = 6).

| Comm.<br>G+C | # Labeled<br>genomes<br>per<br>sample | Buoyant<br>Density<br>window<br>g/ml | $\tilde{\Delta}$ Read<br>coverage<br>X<br>(p-value) | $\tilde{\Delta}$ Proportion<br>of genome<br>recovered in<br>reads<br>(p-value) | $\tilde{\Delta}$ Proportion<br>of genome<br>recovered in<br>contigs<br>(p-value) | $\tilde{\Delta}$ NGA50<br>bp<br>(p-value) | $\tilde{\Delta}$ Proportion<br>of genome<br>recovered<br>in best bin<br>(p-value) | $\tilde{\Delta}$ Proportion<br>bin<br>contaminated<br>(p-value) |
| --- | --- | --- | --- | --- | --- | --- | --- | --- |
| Low | 25 | 1.72-1.77 | 2.202<br>($< 0.001$) | 0.445<br>($< 0.001$) | 0.058<br>(0.027) | 18409<br>($< 0.001$) | 0.675<br>($< 0.001$) | -0.007<br>(NS) |
| | 50 | | 1.769<br>($< 0.001$) | 0.391<br>($< 0.001$) | 0.052<br>($< 0.001$) | 19672<br>($< 0.001$) | 0.567<br>($< 0.001$) | -0.038<br>($< 0.001$) |
| | 100 | | 1.188<br>($< 0.001$) | 0.324<br>($< 0.001$) | 0.006<br>(NS) | 7779<br>($< 0.001$) | 0.337<br>($< 0.001$) | -0.010<br>($< 0.001$) |
| High | 50 | 1.70-1.75 | 0.018<br>(NS) | 0.013<br>(NS) | -0.032<br>(NS) | -212<br>(NS) | 0.019<br>(NS) | -0.001<br>(NS) |
| | | 1.72-1.77 | 0.589<br>($< 0.001$) | 0.241<br>($< 0.001$) | 0.026<br>(NS) | 3824<br>($< 0.001$) | 0.330<br>($< 0.001$) | -0.017<br>(0.006) |
| | | 1.75-1.79 | 3.935<br>($< 0.001$) | 0.388<br>($< 0.001$) | 0.102<br>(0.013) | 47027<br>($< 0.001$) | 0.825<br>($< 0.001$) | -0.012<br>(NS) |

### Metagenomic-SIP improves assembly of isotopically labeled genomes

We found that metagenomic-SIP improved assembly of isotopically labeled genomes over conventional shotgun metagenomics. When sequenced at relatively low depth, metagenomic-SIP allowed for a significantly greater proportion of each labeled genome to be assembled compared to the shotgun metagenomes in all three reference sets (Fig. 3A, Table 2). At high sequencing depth metagenomic-SIP improved target genome assembly in the low and medium G+C reference sets but not in the high G+C set. In all simulations, metagenomic-SIP improved quality of assembled contigs from labeled genomes, as measured by NGA50 (Fig. 3C, Table 3), and more incorporators were assembled at ≥ 50% completeness compared to the shotgun method (Table S3). Considering the need to have at least 50% of the genome recovered to calculate the NGA50 from both metagenomic-SIP and shotgun metagenomic assemblies, our NGA50 analysis was limited to a subset of relatively well assembled references. In all cases, but most notably with the high G+C reference set, incorporators with higher G+C were recovered to a higher proportion by assembled contigs with metagenomic-SIP over shotgun metagenomics (Fig. S11). Similarly, assembly improvement with metagenomic-SIP was greatest for low abundance incorporators, a trend most prominent with the lower G+C genome set (Fig. S11 and S12).

**Fig. 3:**
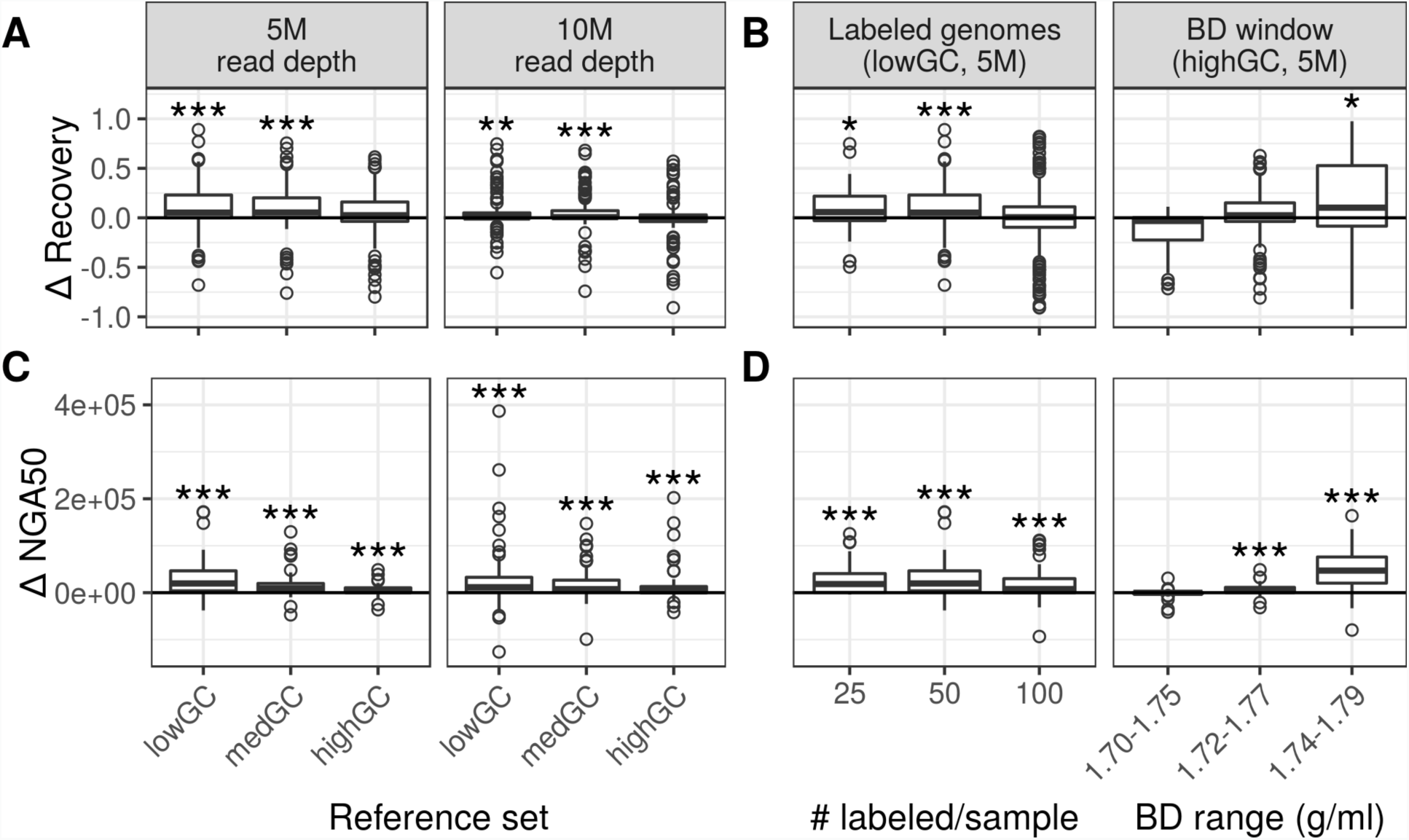
Improvement of metagenomic-SIP relative to conventional metagenomics with respect to co-assembly quality of ^13^C-labeled genomes. Values greater than zero indicate improvement of metagenomic-SIP relative to conventional metagenomics (Bonferroni correction, *n* = 6; *, *p* - values < 0.05; **, *p*-values < 0.01; ***; *p*-values < 0.001). A and C) The difference in proportion of each labeled genome (*n* = 100) recovered (*i.e.* aligned to contigs) and the difference in their NGA50 (in base pairs; where > 50% of the genome is aligned to contigs in both metagenomic simulation types), respectively, varies between communities that differ in G+C content when sequenced at different depths. B and D) The difference in proportion of each labeled genome recovered (*i.e.* aligned to contigs, *n* = 50, 100, 200) and the difference in their NGA50 (in base pairs; where > 50% of the genome is aligned to contigs in both metagenomic simulation types), respectively, for low G+C communities with respect to the number of labeled genomes per sample and for high G+C communities with respect to the position of the BD window.

The number of incorporators per sample influenced how metagenomic-SIP improved assembly of labeled genomes over shotgun metagenomics. Overall, both genome recovery in assembled contigs and NGA50 showed greater improvement with metagenomic-SIP when fewer genomes were labeled per sample. Metagenomic-SIP did not significantly increase the proportion of incorporator genomes recovered in contigs when we labeled 100 genomes per sample (Fig. 3B and D, Table 4). The BD analysis window greatly impacted recovery and quality of contigs in metagenomic-SIP. Metagenomic-SIP improved assembly the most for the heaviest BD window (1.74-1.79 g/ml; Fig. 3B and D, Table 4). The benefit of metagenomic-SIP over shotgun metagenomics was strongly influenced by the G+C of the labeled genome interacting with the BD window. For high G+C incorporators, using a light BD window (1.70-1.75 g/ml) did little to improve assembly over shotgun metagenomics while for the low G+C incorporators we saw little assembly improvement when using a heavy window (1.74-1.79 g/ml; Fig. S13 and S14).

### Metagenomic-SIP improves MAG binning of isotopically labeled genomes

Finally, we found that metagenomic-SIP improved the binning of labeled genomes over conventional shotgun metagenomics. Across all simulations, more labeled genomes were recovered as MAGs with metagenomic-SIP compared to the conventional approach (Table S3). We observed a high accuracy in MAG binning for both metagenomic-SIP and shotgun metagenomic datasets, with a minority of genomes divided or overlapping among multiple bins. Since some genomes were found in multiple bins, we examined binning quality in two ways. First, we used the single “best” bin for each labeled genome (*i.e.* the bin that covered the highest proportion of the reference genome). We found that a greater proportion of each labeled genome was recovered in the best bin when using metagenomic-SIP compared to shotgun metagenomics (Fig. 4A, Table 3). We further found that, for genomes that were successfully binned in both simulation types, metagenomic-SIP bins had less contamination from other genomes compared to the corresponding shotgun metagenomic bins (Fig. 4C, table 2). Second, we combined the multiple bins identified as originating from the same reference genome. This multiple bin approach also showed improved labeled genome recovery and contamination relative to conventional shotgun metagenomics approaches (Fig. S15). Metagenomic-SIP improved binning the most at lower levels of sequencing depth.

**Fig. 4:**
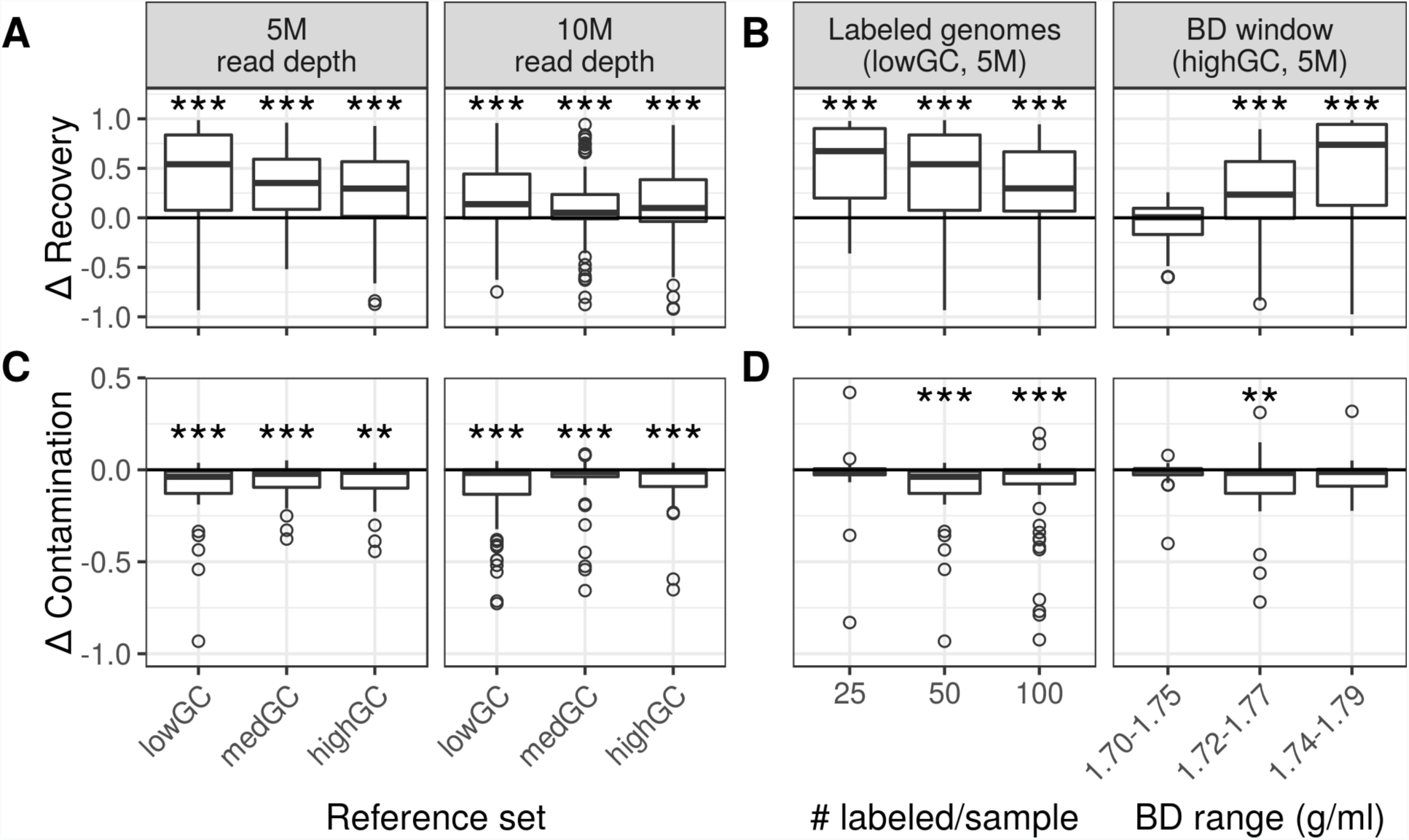
Improvement of metagenomic-SIP relative to conventional metagenomics with respect to MAG bin quality of ^13^C-labeled genomes. For genome recovery in bins, values greater than zero indicate improvement of metagenomic-SIP relative to conventional metagenomics, while for bin contamination values less than zero indicate improvement of metagenomic-SIP relative to conventional metagenomics (Bonferroni correction, *n* = 6; **, *p*-values < 0.01; ***, *p*-values < 0.001). A and C) The difference in the proportion of each labeled genome (*n* = 100) recovered in a bin and the difference in their proportion bin contamination, respectively, varies between communities that differ in G+C content when sequenced at different depths. B and D) The difference in the proportion of each labeled genome (*n* = 50, 100, 200) recovered in a bin and the difference in their proportion bin contamination, respectively, for low G+C communities with respect to the number of target genomes per sample and for high G+C communities with respect to the position of the BD window.

We further found that both the number of incorporators per sample and the choice of the BD window to sequence influenced metagenomic-SIP binning improvement. We tested this using the single best bin approach. With more incorporators per sample, metagenomic-SIP did not improve labeled genome recovery within a bin as much as in simulations with fewer incorporators per sample. Metagenomic-SIP improved bin recovery the most when sequencing a heavier BD window (Fig. 4B, Table 4). Interestingly, moderate values for both number of incorporators per sample (50) and sequenced BD window (1.72-1.77 g/ml) showed the greatest improvement in bin contamination (Fig. 4D, Table 4).

## Discussion

The utility of MetaSIPSim for evaluating metagenomic-SIP efficacy was evident in our comparison of metagenomic-SIP relative to conventional shotgun metagenomics. Metagenomic-SIP improved the ability to assemble and bin isotopically labeled target genomes with higher quality, greater completeness, and less contamination than could be achieved through the application of conventional shotgun metagenomic sequencing. Examples of these improvements for three individual genomes is shown in Fig. 5. Our analyses confirmed that metagenomic-SIP is an effective method for targeted assembly of genomes from complex metagenomes and helped identify cases where the benefits of this method were marginal. Using MetaSIPSim, researchers can use preliminary knowledge of their system’s community composition and the planned experimental parameters to test how well metagenomic-SIP will benefit them and adapt or optimize their sampling, methods, or sequencing regime accordingly.

**Fig. 5:**
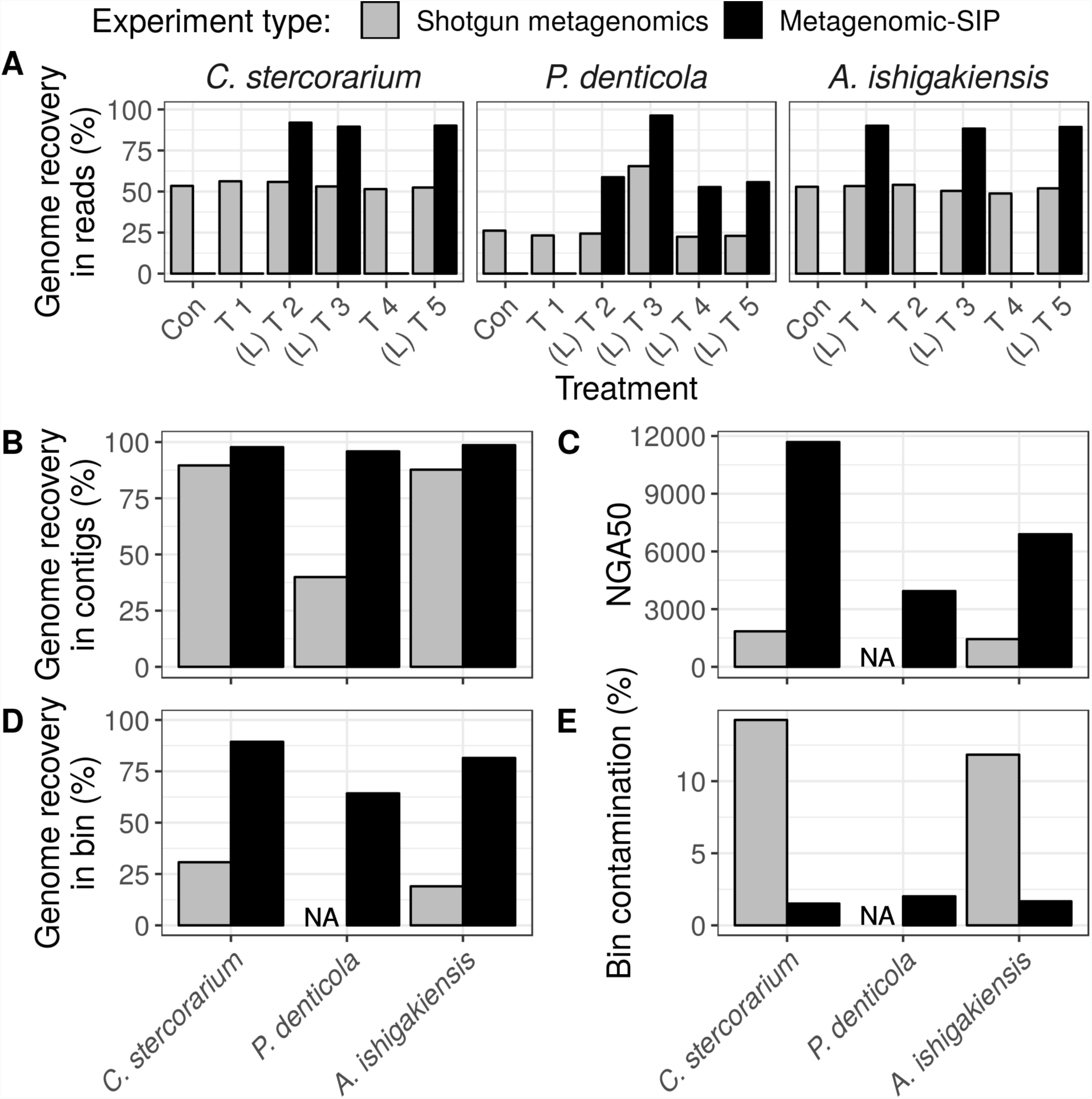
Examples of MAG quality improvements achieved with metagenomic-SIP relative to conventional shotgun metagenomics for three target genomes of low abundance in the community. These examples are taken from the 50% G+C skewed (medGC) reference set sequenced at 5,000,000 reads. Genomes presented here are *Clostridium stercorarium*, *Prevotella denticola*, and *Altererythrobacter ishigakiensis*. A) Percentage of each genome recovered in reads across 6 simulation trials in which community composition and ^13^C-labelling were varied randomly (T1 – T5); ‘con’ indicates the ^12^C-control, ‘L’ indicates a trail in which the organism was ^13^C-labeled. B) Percentage of each genome recovered in contigs from the co-assembly of all 6 trials. C) The NGA50 of the contigs from the co-assembly mapped to each genome. D) The percentage of each genome recovered in a MAG bin. E) The percentage each MAG bin that is contamination from other genomes. Note that we have no NGA50 for the shotgun metagenomics assembly of *P. denticola* as less than 50% of this genome was recovered from the co-assembly. Similarly, we recovered no bin mapping to *P. denticola* from the shotgun metagenome.

Our analyses revealed how experimental design influences the power of metagenomic-SIP. We found that the benefits of metagenomic-SIP relative to conventional shotgun metagenomics declined as sequencing depth and genome coverage increased. We believe that this is largely due to the fact that, with deep sequencing, there is enough read coverage to capture all or most of a community and therefore enrichment with stable isotopes is unnecessary. Our simulations were limited to 500 well characterized bacterial genomes, while natural communities, such as soil, contain orders of magnitude more diversity and complexity [42]. Hence, the utility of metagenomic-SIP is likely to be greater with natural communities relative to the simulations we performed. Further, the ability to achieve greater bin quality with less sequencing coverage may make metagenomic-SIP more appealing for studies that require many samples or replicates due to the trade-off between sequencing depth and sample number [43].

The benefits of metagenomic-SIP varied across bacterial communities that differed in G+C skew. Specifically, we saw that metagenomic-SIP improved assembly and binning of incorporators more in the high G+C skewed communities than in the lower G+C skewed communities. This result is likely due to the incursion of reads from high G+C unlabeled genomes, which are naturally recovered in heavy BD windows. Due to this phenomenon, we recommend a careful selection of the BD window to be sequenced, based on the overall community G+C for metagenomic-SIP studies employing the heavy-SIP method. We confirmed this result by simulating reads from a high G+C genome set with both a lighter and heavier BD window than originally tested. We found that for a community with overall high G+C skew, a heavier BD window resulted in the greatest benefit from metagenomic-SIP.

Regardless of overall community, the benefits of metagenomic-SIP in recovery of an isotopically labeled genome depends on the individual incorporator’s G+C. Specifically, metagenomic-SIP was less advantageous for recovering with low G+C incorporators than for medium to high G+C incorporators. Low G+C genomes are too light to sufficiently shift into the BD window that is sequenced, despite isotopic labeling. This loss of low G+C genomes may be more extreme if heavier or narrower BD ranges are chosen. Indeed, we observed this outcome with our follow-up simulation when using a heavier BD window (Fig. S10 and S16). If a study is designed to specifically target low G+C genomes, such as some *Firmicutes* species, a lighter BD window may be optimal for MAG recovery.

Finally, the benefits of metagenomic-SIP were greatest for incorporators present in low abundance in the community. Most highly abundant incorporators had high-quality assemblies and bins with both metagenomic methods, yet metagenomic-SIP greatly improved assembly and binning over conventional shotgun metagenomics for lesser abundant incorporators. We conclude that metagenomic-SIP shows great promise for metagenome recovery of very low abundant genomes [22] and have shown examples of this potential with three genomes simulated at low relative abundance (Fig. 5).

For our case study, we simulated a multi-substrate SIP experiment. However, there are a number of other experimental designs popularly used dependent on available resources and study hypotheses. One commonly used simple design is to add a single labeled substrate to single or replicate environmental samples. The goal of this design is to use the isotopic labeling to selectively enrich for labeled genomes and may be useful for identifying slow growing, low abundant organisms that utilize a specific substrate. We predict that this method will perform similarly to the multi-substrate method that we simulated and if sequenced to the same depth, may allow for recovery of even more reads from the target genomes. However, without the variation in coverage due to differential labeling across multiple treatments, assembly and MAG binning may produce poorer results than shown here. Another common metagenomic-SIP experimental design is to use a single isotopically labeled substrate in a time series. This design aims to identify successive changes in active populations utilizing a given substrate and to identify trophic networks. Simulations with this experimental design would be very similar to our analyses. The primary difference being that the user may want to include additional unlabeled control samples, one per timepoint. We have not tested either of these designs here but we encourage using MetaSIPSim to test the benefits of these methods over conventional metagenomics for unique experimental designs and parameters.

In addition to testing the benefits of metagenomic-SIP, MetaSIPSim can be used in lieu of *in vitro* metagenomic-SIP experiments with mock communities to generate datasets for development of analytical pipelines, saving time and money. Datasets generated *in silico* with MetaSIPSim can be used to develop and test tools and pipelines specialized for assembly and binning isotopically labeled genomes. Most current metagenomic-SIP studies utilize a heavy window methodology. While this method may currently be the best option in most cases, with advancements to sequencing technologies and high-throughput methodologies, other methods similar to HR-SIP [2] may be practical, utilizing multiple sequenced gradient fractions. HR-SIP-like methods may be useful for overcoming some of the previously described factors that interfere with metagenomic-SIP genome recovery such as community or incorporator G+C but require new analytical tools. Simulated datasets are especially important for development of methods used to identify isotopically labeled contigs or MAGs. By having known reference genomes, atom % excess values for each incorporator, and community profiles, developers can measure sensitivity and specificity of their tools. Further, simulations generated with MetaSIPSim are reproducible, allowing for comparisons between analysis tools.

We can also see MetaSIPSim or a derivative being incorporated into a metagenomic-SIP analysis pipeline to identify labeled contigs, MAGs, or genomes. In such an approach, contigs assembled from metagenomic-SIP might be identified as either isotopically labeled or unlabeled by comparing their empirical coverage distributions across a CsCl gradient to simulated distributions produced with metaSIPSim. Incorporating simulated and actual read distributions in such analyses might provide an approach for identifying isotopically labeled DNA directly from metagenomics-SIP experiments. Further, it may be possible to estimate BD shifts of contigs or MAGs based on theoretical fragment or read distributions generated with MetaSIPSim, thereby enabling quantification of isotopic enrichment.

## Conclusions

MetaSIPSim is a useful tool for simulation-based testing of metagenomic-SIP methods. Using MetaSIPSim we found that metagenomic-SIP experiments significantly improve assembly and binning of targeted, isotopically labeled genomes. Further, experimental design of metagenomic-SIP experiments influences the benefits of this method. We believe that MetaSIPSim can be used to optimize experimental parameters and test new analytical techniques to advance research using metagenomic-SIP.

## Availability and requirements

Project name: MetaSIPSim

Project homepage: https://github.com/seb369/MetaSIPSim

Operating system: Linux and Mac OSX

Programming language: Python 2.7

Other requirements: See table S1

License: MIT license

Any restrictions to use by non-academics: None

## List of abbreviations

SIP: Stable isotope probing BD: Buoyant density
MAG: Metagenome assembled genome
HR-SIP: High-resolution stable isotope probing DBL: Diffusive boundary layer

## Declarations

### Ethics approval and consent to participate

Not applicable

### Consent for publication

Not applicable

### Availability of data and materials

All reference genomes used in these simulations are available through the NCBI Reference Sequence Database (https://www.ncbi.nlm.nih.gov/refseq/). Code used to generate simulations are available at https://github.com/seb369/MetaSIPSim.

### Author contributions

DHB conceived of the project, SEB designed and wrote the software and conducted the analysis, SEB wrote the manuscript with input and edits from DHB.

## Acknowledgements

We thank Nicholas Youngblut for advice and suggestions during development of MetaSIPSim as well as Roland Wilhelm for discussions and review of this manuscript.

