## Supplemental material for "Simulating metagenomic stable isotope probing datasets with MetaSIPSim"

### Supplementary Tables

**Table S1:** MetaSIPSim dependencies

| **Python module/dependency** | **version** |
|---|---|
| numpy | ≥ 1.16.3 |
| pandas | ≥ 0.24.2 |
| Biopython | ≥ 1.73 |
| scipy | ≥ 1.2.1 |
| pyfasta | ≥ 0.5.2 |
| InSilicoSeq (recommended) | ≥ 1.3.6 |

**Table S2:** Parameters used for the MetaSIPSim validation and case study simulations. * Values may differ between simulations as described in the materials and methods.

| Parameter | Lueders et al. 2004 | Buckley et al. 2007 | Wawrik et al. 2009 | Case study simulations |
|---|---|---|---|---|
| BD window or fractions | fraction | | | window |
| Simulation endpoint | fragment list | | | read sequences |
| Sequencing window min BD (g/ml) | NA | | | 1.72* |
| Sequencing window max BD (g/ml) | | | | 1.77* |
| Genome coverage with fragments | 100X | | | |
| Fragment length distribution | Skewed normal (mean = 9000, scale = 2500, shape = -5) | | | |
| Temperature (K) | 293.15 | | | |
| Average gradient density (g/ml) | 1.725 | 1.69 | 1.701 | 1.69 |
| (angular velocity)$^2$ (rad/s)$^2$ | 20465612 | 33172837 | 16860743 | 33172837 |
| Min radius of tube from axis of rotation (cm) | 7.21 | 2.6 | 7.47 | 2.6 |
| Max radius of tube from axis of rotation (cm) | 8.49 | 4.85 | 8.79 | 4.85 |
| Angle of tube (degrees) | 0 | 28.6 | 0 | 28.6 |
| Tube radius (cm) | 0.65 | 0.66 | 0.66 | 0.66 |
| Tube height (cm) | 6.6 | 4.7 | 4.7 | 4.7 |
| Proportion of DNA in DBL | 0.001 | | | |
| Stable isotope element | C | N | N | C |
| Min BD for model gradient (g/ml) | 1.67 | | | 1.67* |
| Max BD for model gradient (g/ml) | 1.775 | | | 1.775* |
| BD steps for model gradient (g/ml) | 0.0001 | | | 0.0001 |
| Max read length (bp) | NA | | | 151 |
| Average insert size (bp) | | | | 1000 |
| Standard deviation of insert size (bp) | | | | 5 |
| Final number of reads | | | | 5,000,000* |

**Table S3:** Summary statistics for recovery of reads, contigs, and MAGs for labeled genomes in initial simulations.

| Metagenome type | Comm. G+C | Seq. depth | Genomes aligned to contigs (≥ 50% recovered) | Genomes aligned to contigs (≥ 90% recovered) | Genomes recovered as MAGs |
|---|---|---|---|---|---|
| SIP | Low | 5MM | 74 | 53 | 64 |
| shotgun | | | 66 | 36 | 32 |
| SIP | Medium | | 66 | 48 | 56 |
| shotgun | | | 61 | 33 | 36 |
| SIP | High | | 59 | 41 | 45 |
| shotgun | | | 57 | 30 | 38 |
| SIP | Low | 10MM | 93 | 75 | 89 |
| shotgun | | | 85 | 69 | 69 |
| SIP | Medium | | 87 | 68 | 73 |
| shotgun | | | 79 | 63 | 67 |
| SIP | High | | 80 | 59 | 67 |
| shotgun | | | 78 | 60 | 56 |

**Supplementary Figures**

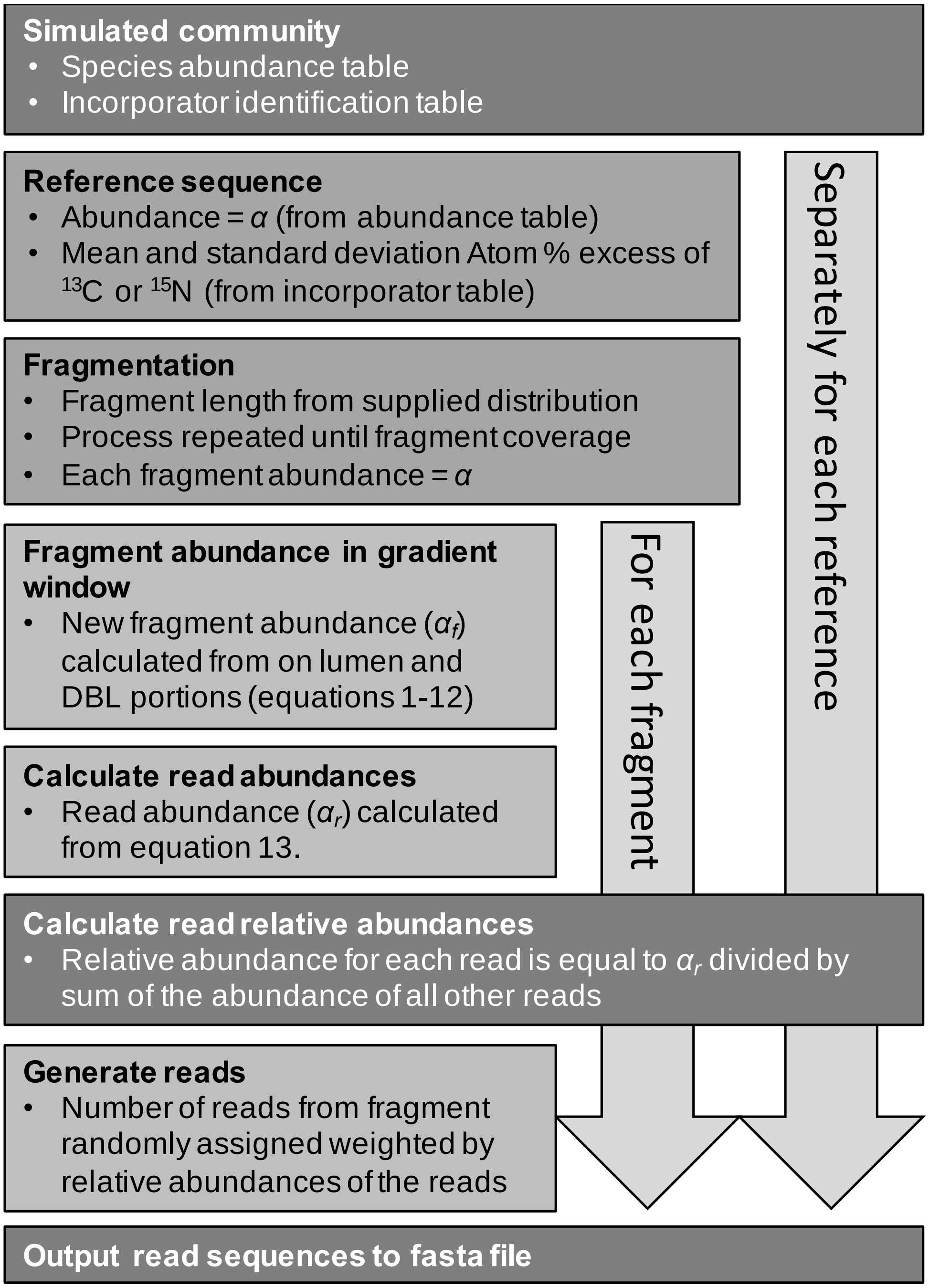


**Fig. S1:** Diagram of the simulation procedure.

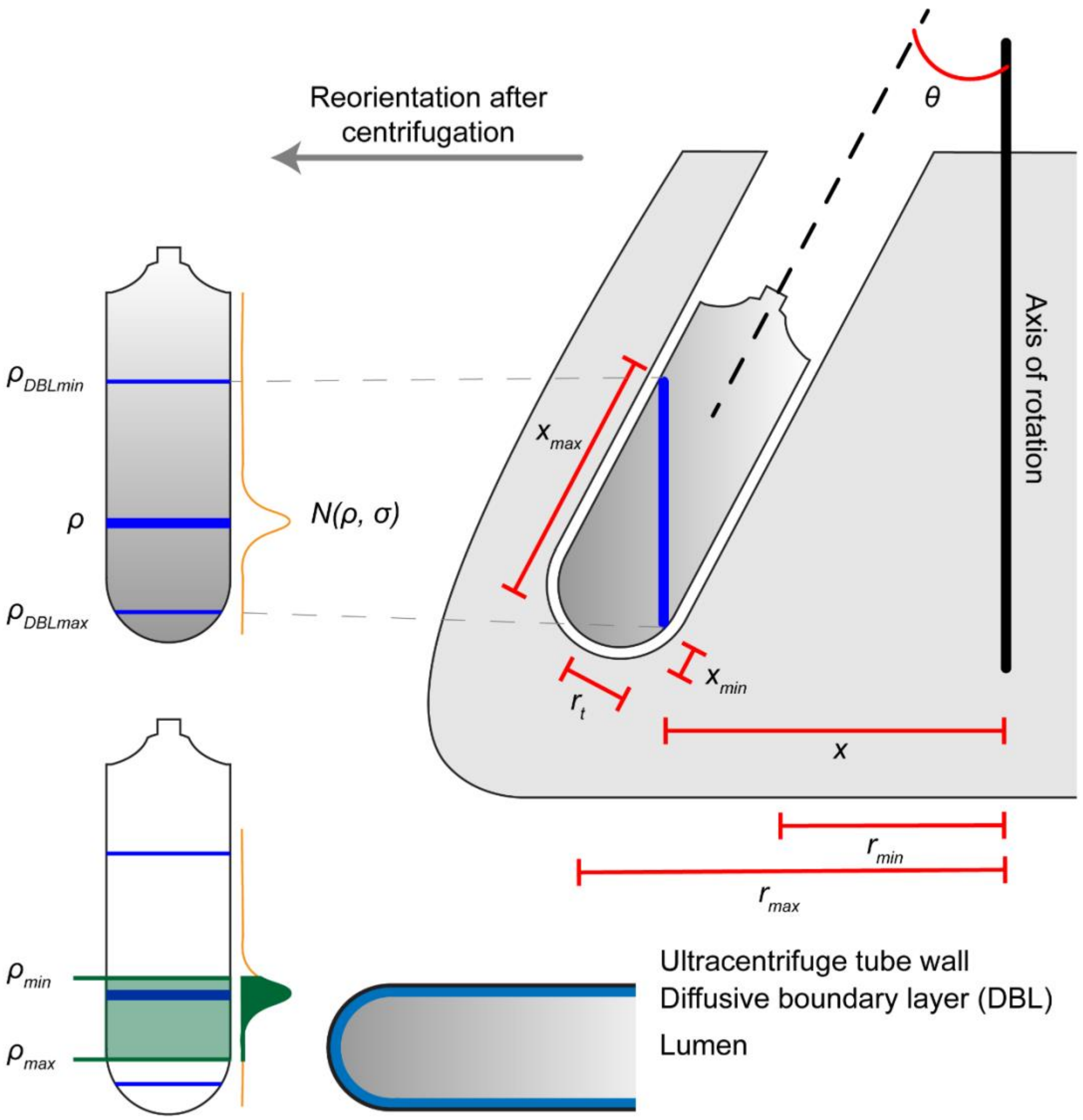


**Fig. S2:** Visual description of variables defined in the manuscript and table 2. Thick blue line indicates the position of a single DNA fragment during and after ultracentrifugation. Centrifugation setup shown here is based on a fixed angled rotor.

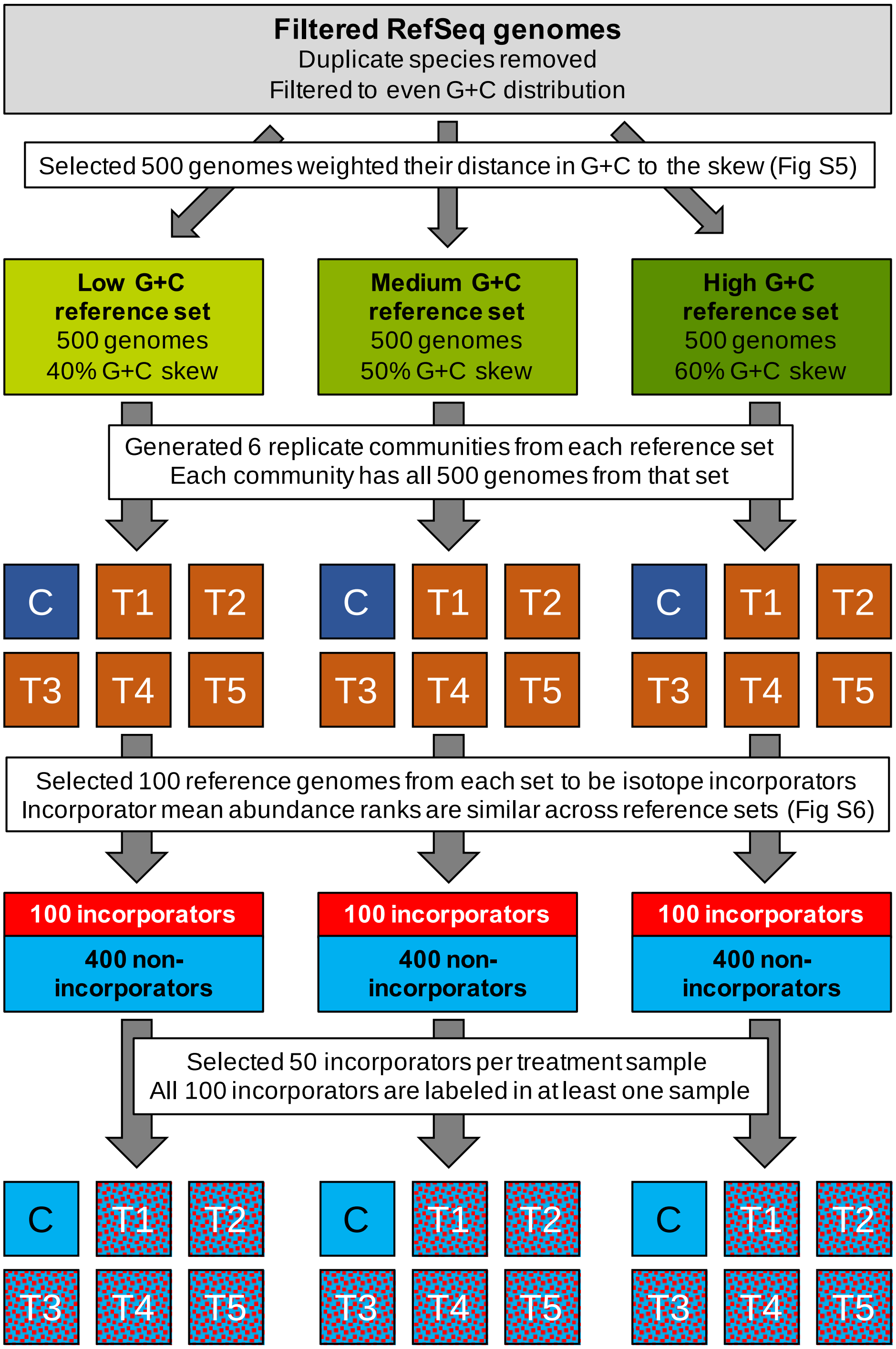


**Fig. S3:** Diagram of experimental design for case study simulations.

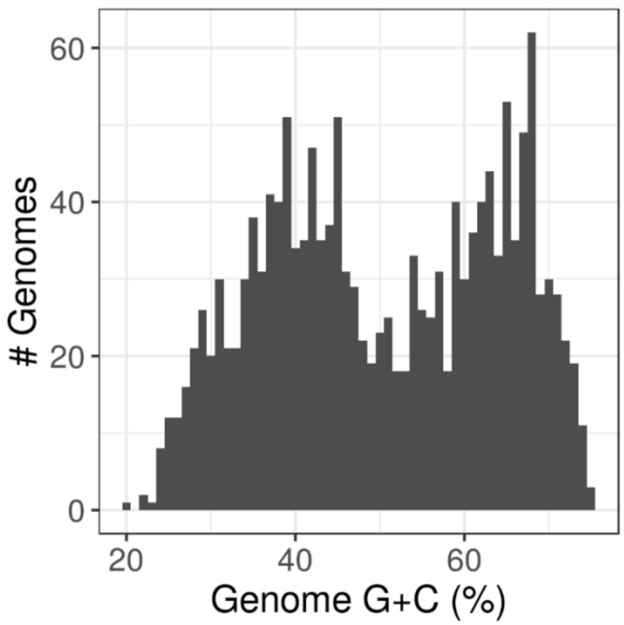


**Fig. S4:** RefSeq genomes G+C distribution with G+C bins rounded to nearest whole number (downloaded January 25, 2019).

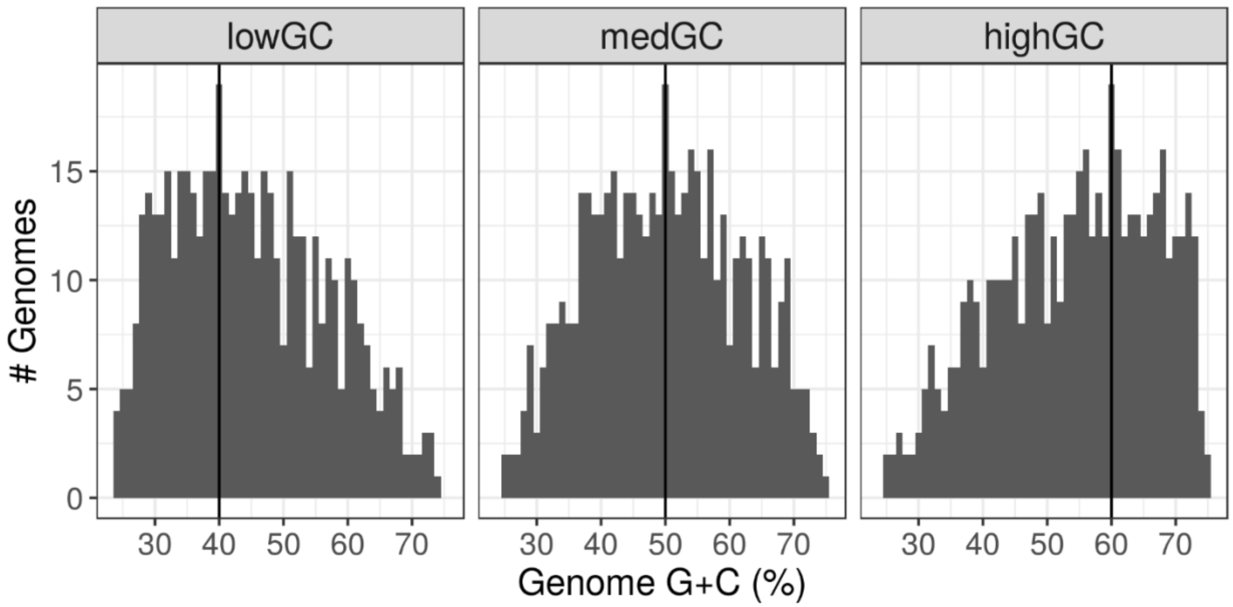


**Fig. S5:** G+C distributions for the 500 genomes in each reference set. G+C bins are rounded to the nearest whole number. Vertical line indicates G+C skew of the set.

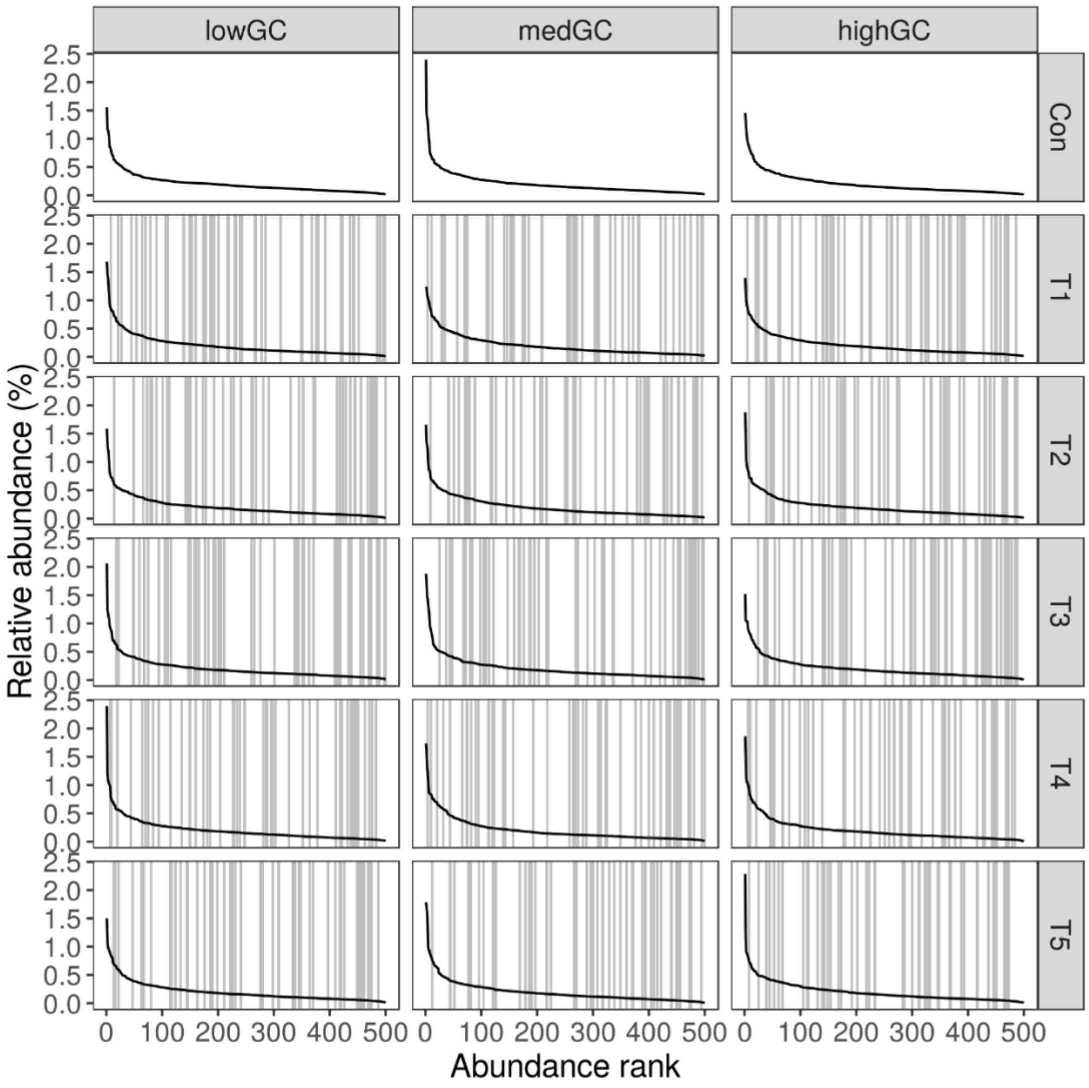


**Fig. S6:** Rank-abundance plots for each simulated sample or replicate community including the $^{12}C$-control (Con) and all five $^{13}C$-trials (T1-T5). Random genomes are only labeled in the $^{13}C$-labeled samples. Vertical grey lines indicate ranks of the 50 labeled genomes per treatment. Labeled genomes were selected such that mean ranks of the labeled genomes averaged across all samples were similar across the reference sets (lowGC, medGC, and highGC).

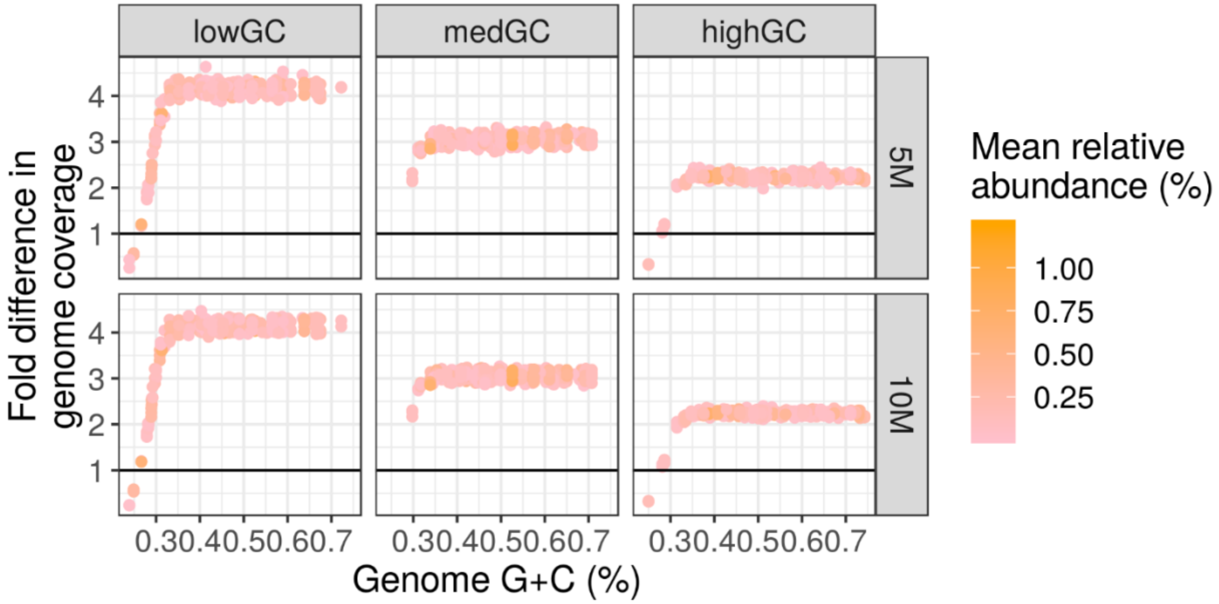


**Fig. S7:** Fold difference in raw read coverage for each labeled genome between the metagenomic-SIP and shotgun metagenomic libraries from the original simulations. Values above one indicate greater coverage in the metagenomic-SIP compared to the shotgun metagenomic libraries.

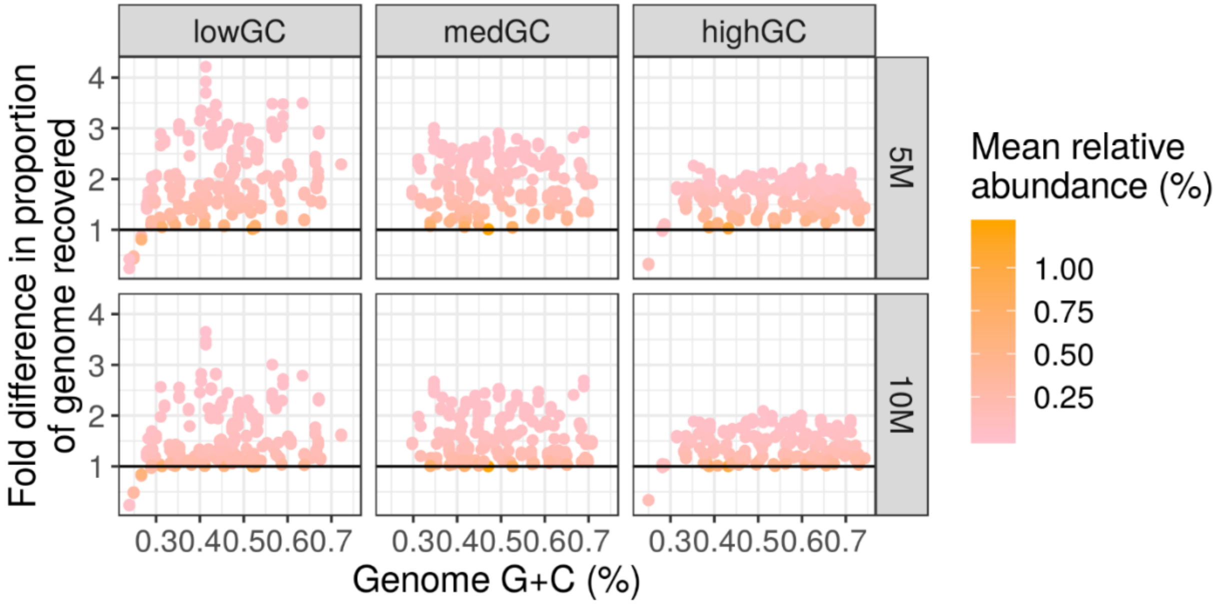


**Fig. S8:** Fold difference in proportion of each labeled genome recovered by reads between the metagenomic-SIP and shotgun metagenomic libraries from the original simulations. Values above one indicate greater recovery in the metagenomic-SIP compared to the shotgun metagenomic libraries.

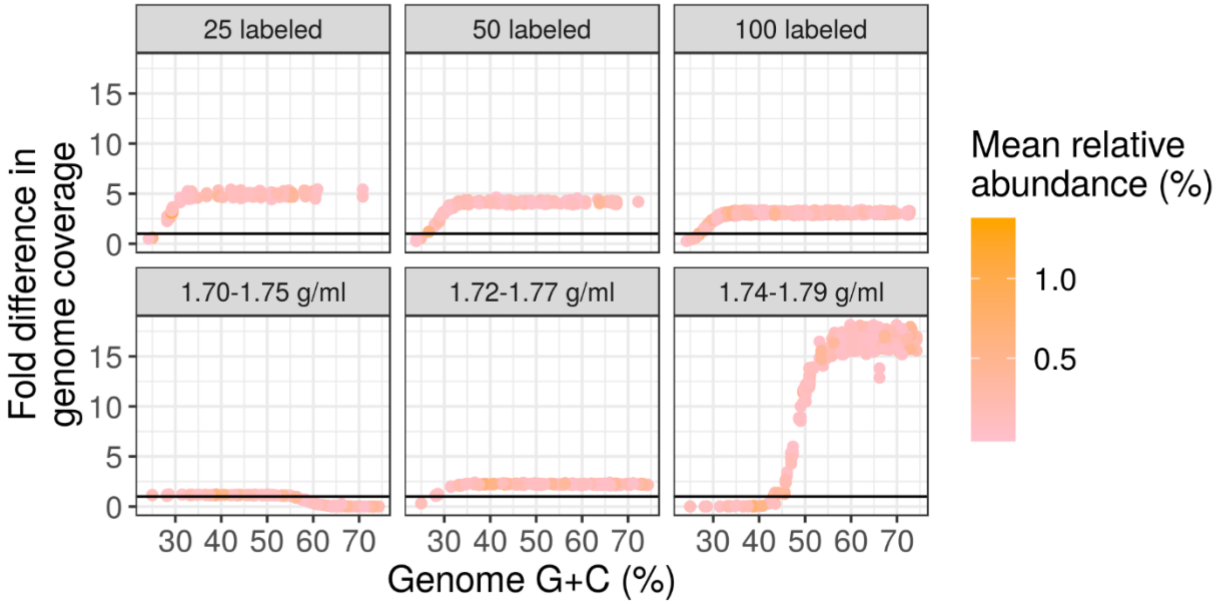


**Fig. S9:** Fold difference in raw read coverage for each labeled genome between the metagenomic-SIP and shotgun metagenomic libraries from the follow-up simulations. Values above one indicate greater coverage in the metagenomic-SIP compared to the shotgun metagenomic libraries. Simulation with the lowGC reference set with varying number of labeled genomes per sample is in the top row while the simulation with the highGC reference set with different sequencing window BD ranges is in the bottom row.

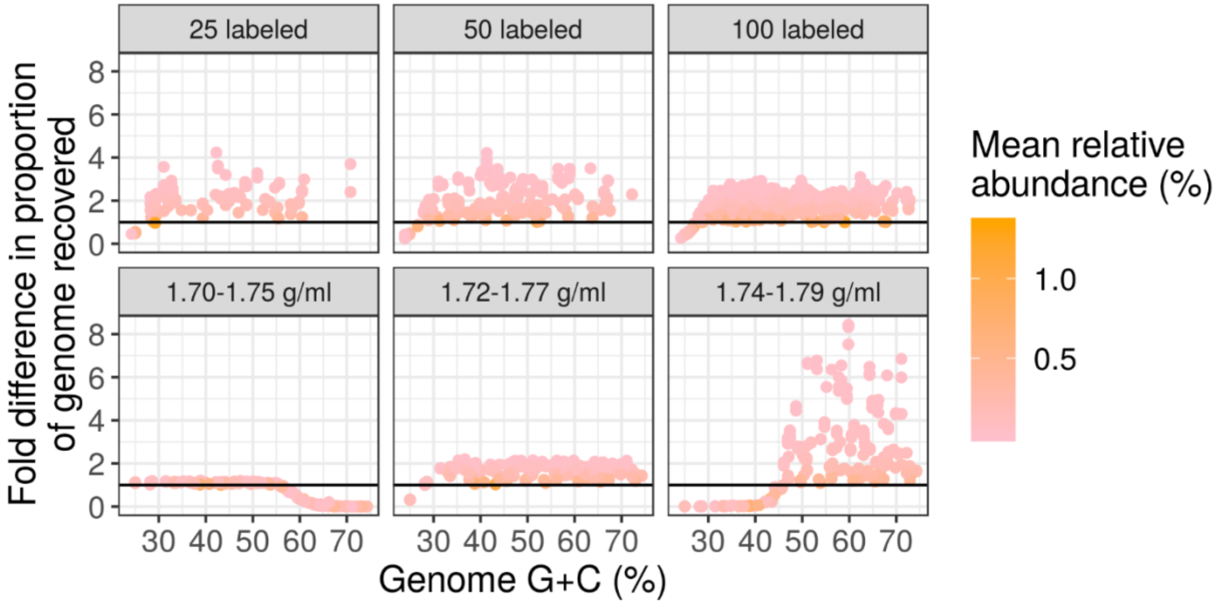


**Fig. S10:** Fold difference in proportion of each labeled genome recovered by reads between the metagenomic-SIP and shotgun metagenomic libraries from the follow-up simulations. Values above one indicate greater recovery in the metagenomic-SIP compared to the shotgun metagenomic libraries. Simulation with the lowGC reference set with varying number of labeled genomes per sample is in the top row while the simulation with the highGC reference set with different sequencing window BD ranges is in the bottom row.

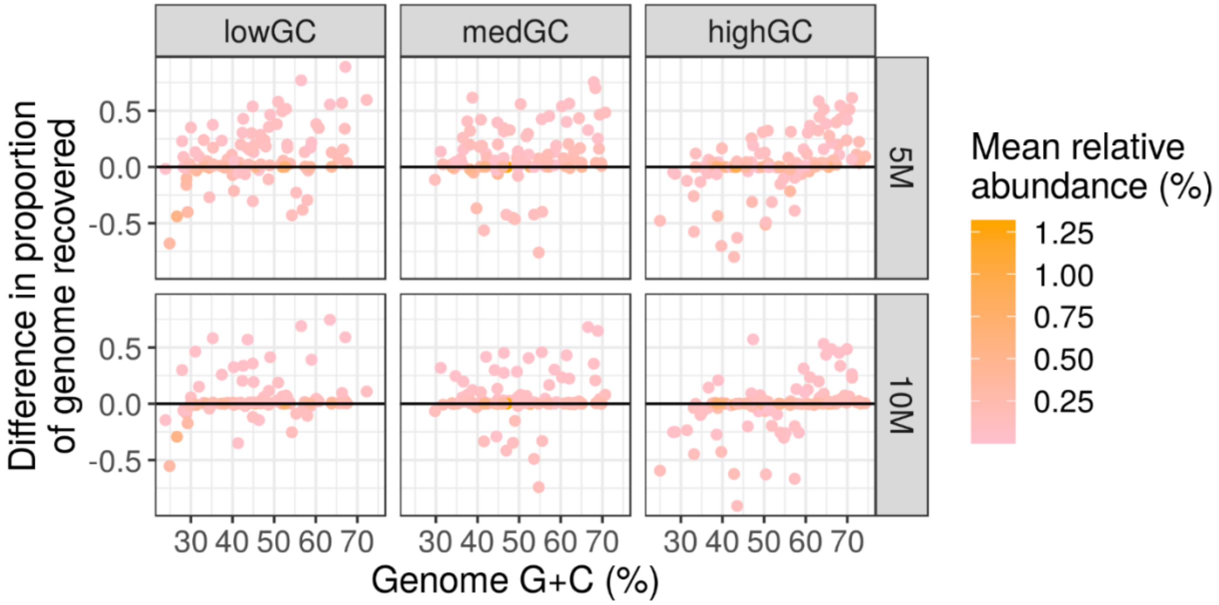


**Fig. S11:** Difference in proportion of each labeled genome recovered in co-assembled contigs between the metagenomic-SIP and shotgun metagenomic libraries from the original simulations. Values above zero indicate greater recovery in the metagenomic-SIP compared to the shotgun metagenomic contigs.

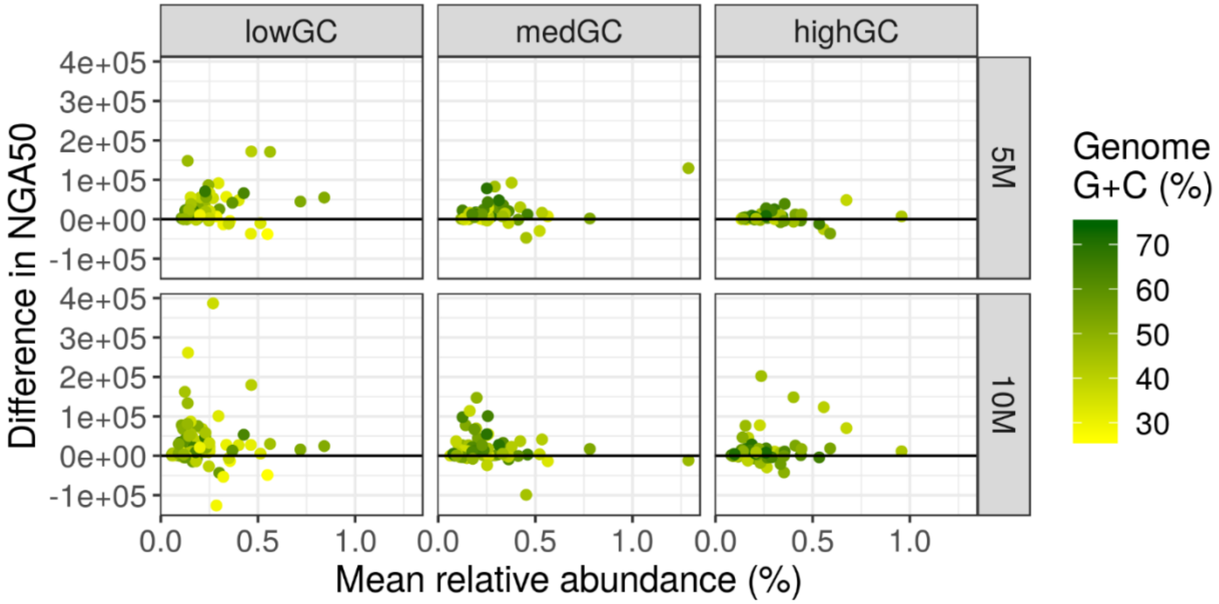


**Fig. S12:** Difference in NGA50 of each labeled genome covered by co-assembled contigs between the metagenomic-SIP and shotgun metagenomic libraries from the original simulations. Values above zero indicate greater NGA50 in the metagenomic-SIP compared to the shotgun metagenomic contigs. Only genomes with over 50% recovery in both SIP and shotgun metagenomes were used in this analysis.

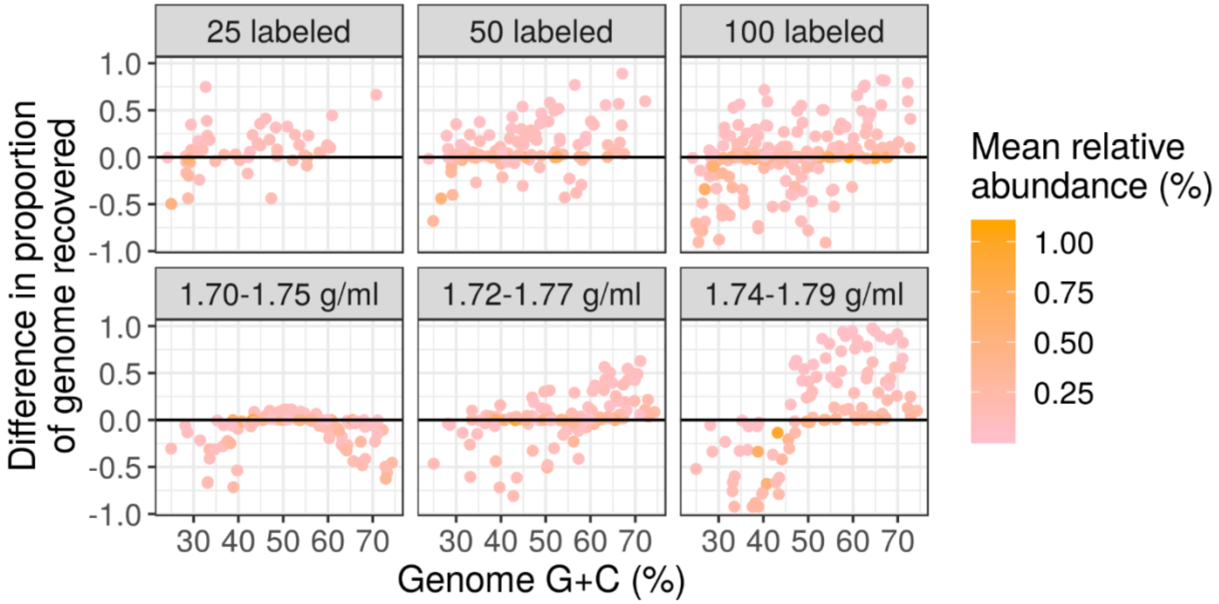


**Fig. S13:** Difference in proportion of each labeled genome recovered in co-assembled contigs between the metagenomic-SIP and shotgun metagenomic libraries from the follow-up simulations. Values above zero indicate greater recovery in the metagenomic-SIP compared to the shotgun metagenomic contigs. Simulation with the lowGC reference set with varying number of labeled genomes per sample is in the top row while the simulation with the highGC reference set with different sequencing window BD ranges is in the bottom row.

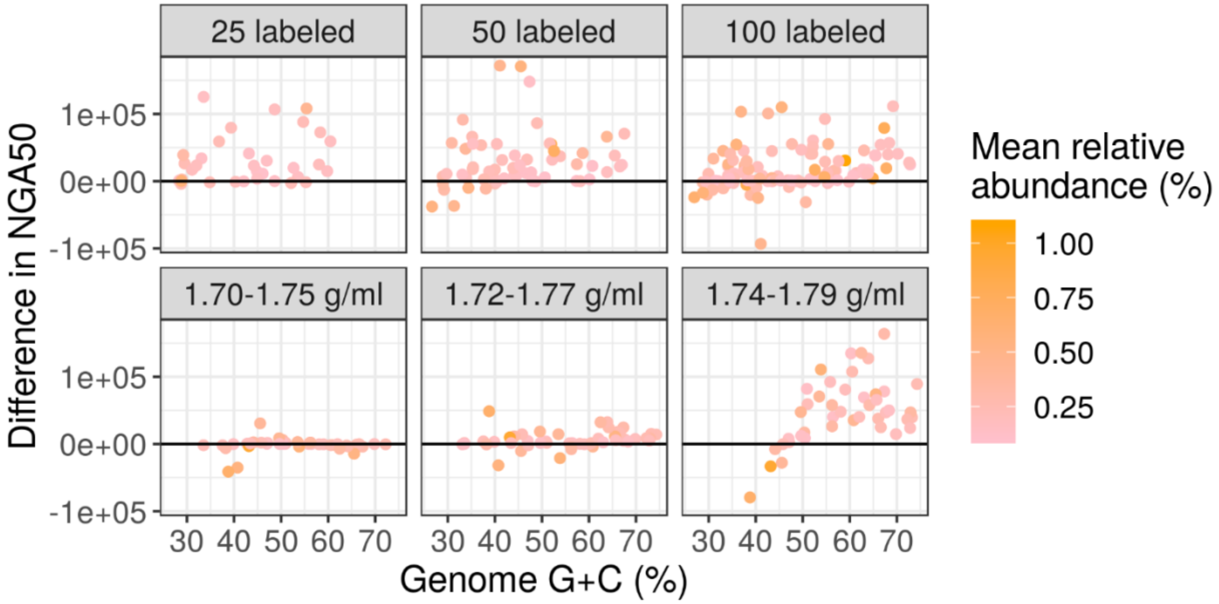


**Fig. S14:** Difference in NGA50 of each labeled genome covered by co-assembled contigs between the metagenomic-SIP and shotgun metagenomic libraries from the original simulations. Values above zero indicate greater NGA50 in the metagenomic-SIP compared to the shotgun metagenomic contigs. Only genomes with over 50% recovery in both SIP and shotgun metagenomes were used in this analysis. Simulation with the lowGC reference set with varying number of labeled genomes per sample is in the top row while the simulation with the highGC reference set with different sequencing window BD ranges is in the bottom row.

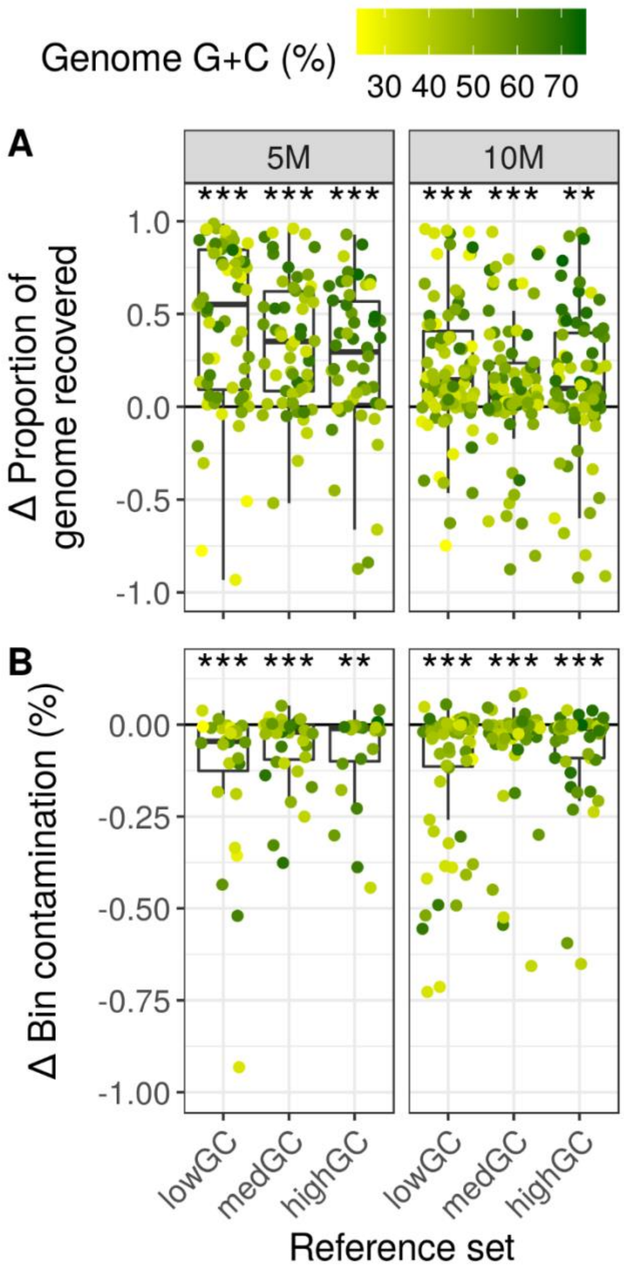


**Fig. S15:** Difference in binning quality between the SIP-metagenomes and shotgun metagenomes with the initial simulations using multiple bins per labeled genome. A) Difference in proportion of each labeled genome recovered in bins. B) Difference in the cumulative contamination for each labeled genome bin set.

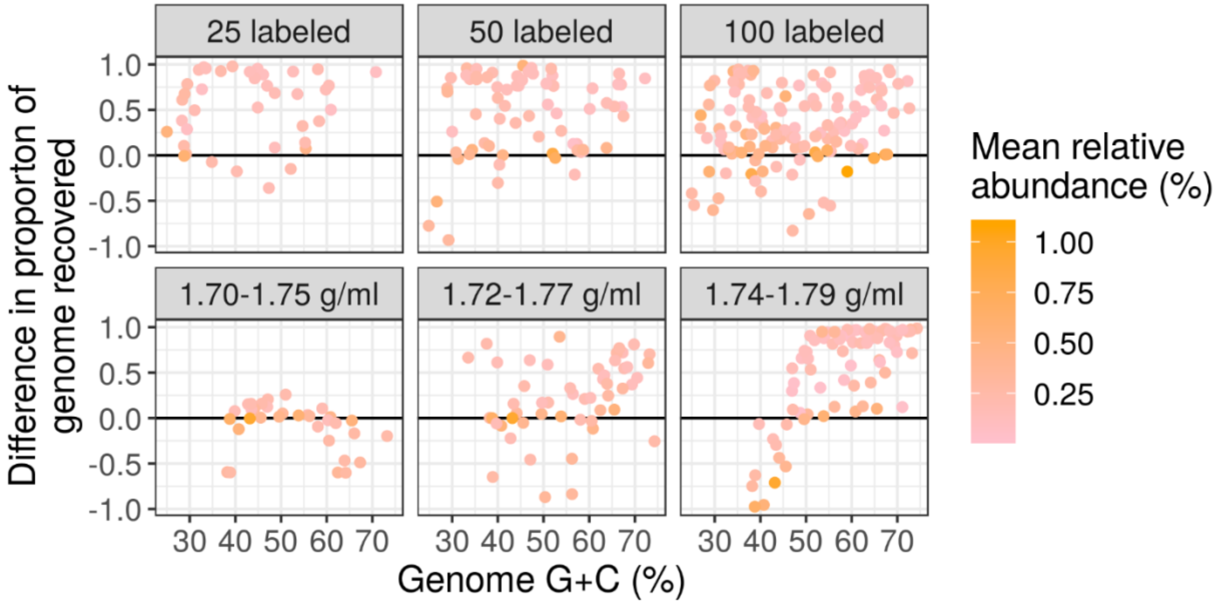


**Fig. S16:** Difference in proportion of each labeled genome recovered in a single most complete bin between the metagenomic-SIP and shotgun metagenomic libraries from the follow-up simulations. Values above zero indicate greater recovery in the metagenomic-SIP compared to the shotgun metagenomic bins. Simulation with the lowGC reference set with varying number of labeled genomes per sample is in the top row while the simulation with the highGC reference set with different sequencing window BD ranges is in the bottom row.
